# Metabolomics-informed coarse-grained model enables prediction of cell-free protein expression dynamics

**DOI:** 10.1101/2025.06.20.660830

**Authors:** William Poole, Manisha Kapasiawala, Ankita Roychoudhury, Matthew Haines, Paul Freemont, Richard M. Murray

**Author notes:** To whom correspondence should be addressed; E-mail: wp. Contributed equally (William Poole, Manisha Kapasiawala).

## Abstract

Cell-free expression systems have gained considerable attention for their potential in biomanufacturing, biosensing, and circuit prototyping. All these endeavors make use of the innate metabolic activity of cell lysates. However, our knowledge of the underlying nature of cell-free metabolism remains lacking. In this work, we use non-targeted mass spectrometry to generate time-course measurements of small molecules in *E. coli* cell lysates (metabolomics) during cell-free protein synthesis (CFPS) to show that the majority of *E. coli*’s metabolism is active in cell lysate. Furthermore, these data indicate that protein synthesis is a relatively small metabolic burden on cell lysates compared to the background metabolism, and that the build-up of multiple metabolic waste products is fundamentally responsible for stopping cell-free protein expression as opposed to depletion of fuel sources. We use these insights coupled with high-throughput CFPS experiments to develop a foundational coarse-grained mechanistic model of CFPS which we show can be easily recalibrated using Bayesian parameter inference techniques to provide accurate models across hundreds of novel experimental conditions. This work provides new experimental insights into the effects of cell-free metabolism on CFPS and establishes a novel framework for leveraging existing CFPS models in new experimental contexts, opening the avenue to future cell-free applications aimed at building complex systems, from multilayered biological circuits to synthetic biological cells.

## Introduction

Synthetic biology harnesses engineering principles and the versatility of biology to enable the manipulation and creation of biological parts and systems. Among its many tools, cell-free systems have emerged as a resourceful tool with diverse applications [1]. The most widespread use of cell-free systems is cell-free protein synthesis (CFPS), where crude cell lysate or purified proteins are supplied with a DNA template and an energy buffer containing building blocks and co-factors for *in vitro* transcription and translation (TX-TL) [2]. Operating outside living cells, CFPS systems offer many advantages, including the ability to produce biomolecules toxic to cell viability and an open reaction environment compatible with lab automation to accelerate design-build-test-learn cycles. These benefits have enabled applications such as biomanufacturing [3]; rapid prototyping of biological parts, pathways, and circuits [4–8]; point-of-care diagnostics and biosensing [9–11]; elucidating the synthesis and assembly of complex biological structures such as membrane proteins and bacteriophages [12–16]; and providing an avenue for the development of artificial life [17–19]. Crude cell lysate-based systems remain the most popular type of CFPS systems, although the cellular machinery needed to drive CFPS is increasingly provided in the form of a minimal set of proteins necessary and sufficient for CFPS [20, 21]. Beyond TX-TL machinery, lysate-based systems also contain much of the cytoplasmic content of cells – including metabolic enzymes, organelles like the endoplasmic reticulum (in eukaryotic lysates), and inverted membrane vesicles – that support processes such as energy regeneration, membrane protein synthesis, and possibly oxidative phosphorylation [22]. While not directly involved in CFPS, these contents enable high expression of a diverse set of proteins, including those that require post-translational modifications, chaperone proteins, and other requirements beyond simple translation. However, while conferring these advantages, this cytoplasmic content also has many unknown and potentially harmful effects on CFPS that lead to poor predictability of CFPS performance. Improving CFPS predictability is crucial for more advanced applications of lysate-based systems, from integrating synthetic cell subsystems to translating *in vitro* learnings into cellular engineering.

Toward improving the predictability of CFPS systems, many previous studies have focused on understanding and modeling the effects of reaction components, particularly small molecules, on CFPS. One study focused on the effects of molecular crowding agents and magnesium (Mg^2+^) on translation initiation and elongation rates [23]. Another study measured the effects of 20 reaction components on several metrics of CFPS dynamics, including the maximum rate of protein production, the time to reach steady state protein level, and others [24]. A third study focused on characterizing the interactions between CFPS reaction components – specifically, Mg^2+^ and various fuels used to regenerate energy – and their differential effects on total transcription and translation [25].

Recognizing that CFPS variability is derived largely from the uncharacterized contents of cell lysate, some studies have focused more directly on the interaction between CFPS and cell-free metabolism. Varner and co-workers have focused on combining modeling and experimental approaches to create a CFPS model coupled with cell-free metabolism [26, 27]. Using dynamic measurements of nucleotide triphosphates (NTPs), amino acids, and several key metabolites of central carbon metabolism, they were able to construct an ensemble of kinetic models that could predict expression of a reporter protein. Meanwhile, Styczynski and co-workers have used metabolomics to characterize and compare central carbon and amino acid metabolism under various reaction conditions, including the absence/presence of CFPS, targeted supplementation of metabolic enzymes, and different methods of cell lysate preparation [28, 29].

While these papers have made important strides towards understanding the relationship between cell-free metabolism and CFPS, this recent work has focused largely on central carbon, NTP, and amino acid metabolism. The success of cell-free metabolic engineering, which relies partly on using innate cell metabolism to generate biochemical intermediates that are then used as precursors for engineered metabolic pathways, suggests that other parts of cell metabolism may be active in cell-free systems [30]. However, the extent to which all metabolic pathways are active in a cell-free context is unknown. Beyond a broad characterization of cell-free metabolism, integrating knowledge of metabolism into predictive models of cell-free protein synthesis, particularly for a wide range of experimental conditions, remains a challenge.

In this work we used untargeted metabolomics to get a high level view of cell lysate metabolism and found that most *E. coli* metabolic pathways are active in cell lysates, resulting in complex effects on CFPS. These experimental observations were used to motivate a novel coarse-grained mechanistic model of cell lysate metabolism focused on the build-up of metabolic waste products. We fit the model’s parameters using a large CFPS dataset with Bayesian parameter inference techniques and then show that this model can be re-calibrated to accurately predict protein expression dynamics under novel experimental conditions by fine-tuning a subset of its parameters via re-training on smaller condition-specific datasets. We demonstrate that this approach can be applied to predict dynamics under a broad range of new experimental conditions, including various cell lysate batches, DNA concentrations, fuel sources, and more. Finally, we show that the model can be slightly modified to include the effects of Mg^2+^ on CFPS, and that this updated model’s prediction recapitulates a previously observed trade-off between energy regeneration and waste mitigation. This work establishes a novel framework for leveraging existing CFPS models in new experimental contexts and provides new predictive insights into the effects of cell-free metabolism on CFPS.

## Results

### Cell lysate time-course metabolomics suggests *E. coli* metabolism is highly active during CFPS

We used high-throughput untargeted mass spectrometry to broadly identify metabolites [31] which are present and dynamically changing during CFPS in *E. coli* lysate. Data was collected across a diverse set of experimental conditions including duration of the CFPS reaction, different cell culture batches, preparation methods, salt concentrations, and additives. Specifically, we used three different cell culture batches, two different magnesium-glutamate concentrations, the addition or absence of 4 nM of a constitutively-expressing deGFP plasmid, the addition of additional oxygen during the CFPS reactions, and variations in the lysate preparation method (Table 1). The use of many conditions across multiple time points allowed us to statistically find a wide variety of associations in the data. These conditions were distributed in such a way that we could easily pool samples between sets of different conditions and still have enough statistical power to draw meaningful conclusions (see Methods, Table 1). Details of our statistical methodology can be found in the Methods section.

**Table 1:** Cell lysate conditions. For each sample condition, four time points were collected from independent CFPS experiments which were quenched with cold methanol and flash frozen at 0 hours, 3 hours, 6 hours, and 12 hours. This resulted in a total of 48 different metabolomics samples. Around 328 metabolites were identified across all these samples resulting in a total of 15744 individual small molecule measurements. Prep Methods: **FP** = French Press; **D** = Dialysis; **S** = Sonication.

| Condition ID | DNA (nM) | Mg (uM) | Extra O2 | Culture Batch ID | Prep Method |
| --- | --- | --- | --- | --- | --- |
| U1 | 0 | 10 | no | WP2 | FPD |
| U2 | 4 | 10 | no | WP2 | FPD |
| U3 | 4 | 10 | no | WP2 | FPD |
| U4 | 4 | 10 | no | WP2 | FPD |
| U5 | 4 | 5 | no | WP1 | SD |
| U6 | 4 | 5 | no | WP1 | FPD |
| U7 | 4 | 5 | no | WP1 | FP |
| U8 | 0 | 5 | no | WP1 | FPD |
| U9 | 4 | 5 | no | eSC5 | FP |
| U10 | 0 | 5 | no | eSC5 | FP |
| T2 | 4 | 10 | yes | WP2 | FPD |
| T6 | 4 | 5 | yes | WP1 | FPD |

First, we examined how the distributions of individual metabolites change over time by pooling samples by timepoint across all experimental conditions. This analysis showed that 87 of 328 metabolites vary statistically significantly over time (Figure 1A). These metabolites are spread out over a wide swath of *E. coli* metabolism. We then examined if different lysate conditions gave rise to statistical differences in the measured metabolite dynamics. Surprisingly, the vast majority of conditions we compared produced few statistical differences. Notably, the addition of constitutively active deGFP plasmid had no measurable effect on overall lysate metabolism, suggesting that CFPS uses relatively little energy compared to other background metabolic processes. The only factor that we found to have statistically significant differences between individual metabolites was the specific cell lysate batch (Figure 1B), suggesting that variability in growth conditions, when cells are harvested, and other details of the lysate preparation protocol can have significant effects on lysate metabolism.

**Figure 1:**
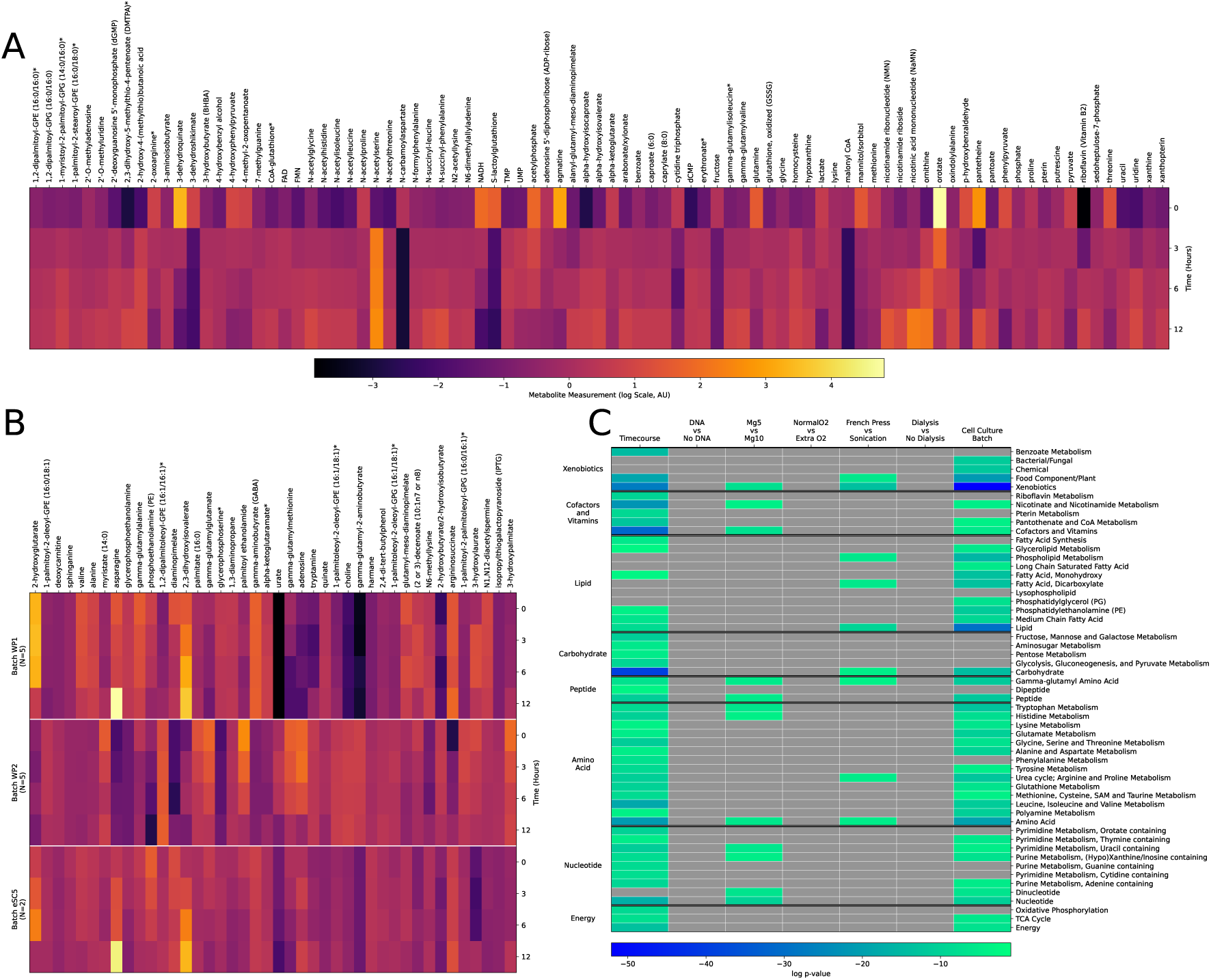
An overview of the significant results from the metabolomics statistical analysis. A. Average time course data for all the metabolites which statistically significantly change in time. B. Average data for individual metabolites which vary between lysate batches. C. Overview of statistically significantly varying pathways.

Next, we simultaneously increased the statistical power of our data and provided a higher-level view of *E. coli* lysate metabolism by statistically combining the measurements of multiple metabolites involved in the same metabolic pathway [32]. This analysis found that 47 out of 55 analyzed pathways vary significantly in time. Cell lysate batch was also revealed to have dramatic changes in 40 pathways, emphasizing our previous findings that growth conditions play a major role in lysate variability. Lysis method (sonication versus French press) and magnesium-glutamate concenentration also resulted in a handful of statistically significant pathway-level metabolic changes (Figure 1C). Strikingly, even with the increased statistical power afforded by pathway level analysis, CFPS did not result in any statistically significant changes in metabolism.

### Fuel and DNA spiking experiments inform a phenomenological model of cell lysate metabolism

We performed a number of additional experiments to understand how cell lysate metabolism affects CFPS and simultaneously understand if CFPS can be used to measure high-level metabolic effects. First, we added a constitutively expressing deGFP construct into cell lysate mixed with energy buffer after various incubation times (Figure 2, middle panel). These experiments show that cell lysate’s innate metabolism diminishes and eventually stops CFPS (as opposed to negative feedback from CFPS itself).

**Figure 2:**
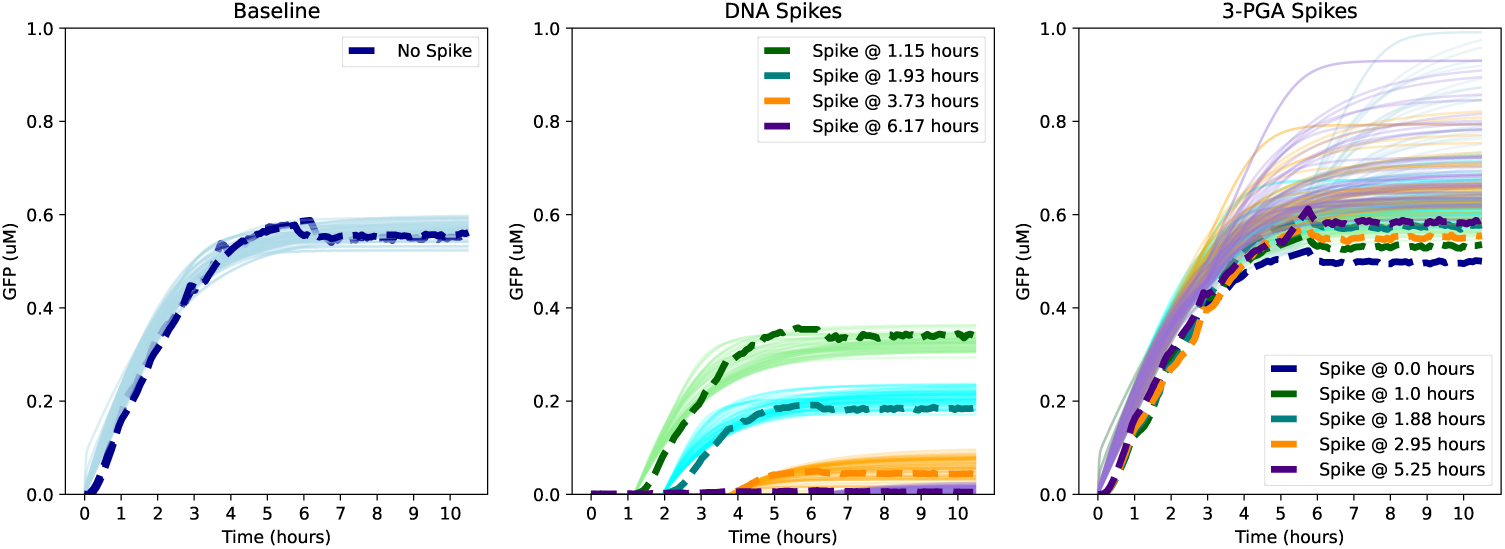
Model simulations for the 100 most likely parameters (light lines) compared to the spiking and baseline data (dashed lines).

In a different set of experiments, we tried adding three additives – additional fuel (3PGA), HEPES buffer, and water – to a CFPS reaction at various times. These results show that adding additional fuel is in fact slightly toxic to cell lysate (Figure 2, right panel; Figure 6). Decreasing pH had been hypothesized to be responsible for loss of cell lysate function, but HEPES buffer does not appear to dramatically increase CFPS (Figure 6). Finally, the water spikes act as a control – it turns out that molecular crowding plays an important role in cell lysate function [33, 34] and therefore relative concentrations and the addition of water can effect CFPS (Figure 6). However, we statistically compared the effects of adding water to those of adding HEPES and 3PGA and found they produced statistically significant differences in the final amount of protein produced. Additionally, when the 3PGA and HEPES experiments are renormalized to account for the effects of adding water, they still have a statistically significant effect effect on CFPS. A breakdown of these statistics can be seen in Table 2. Finally, we measured an ATP time course of cell lysate mixed with energy buffer, which shows that the available amount of ATP diminishes over the course of six hours (Figure 3).

**Figure 3:**
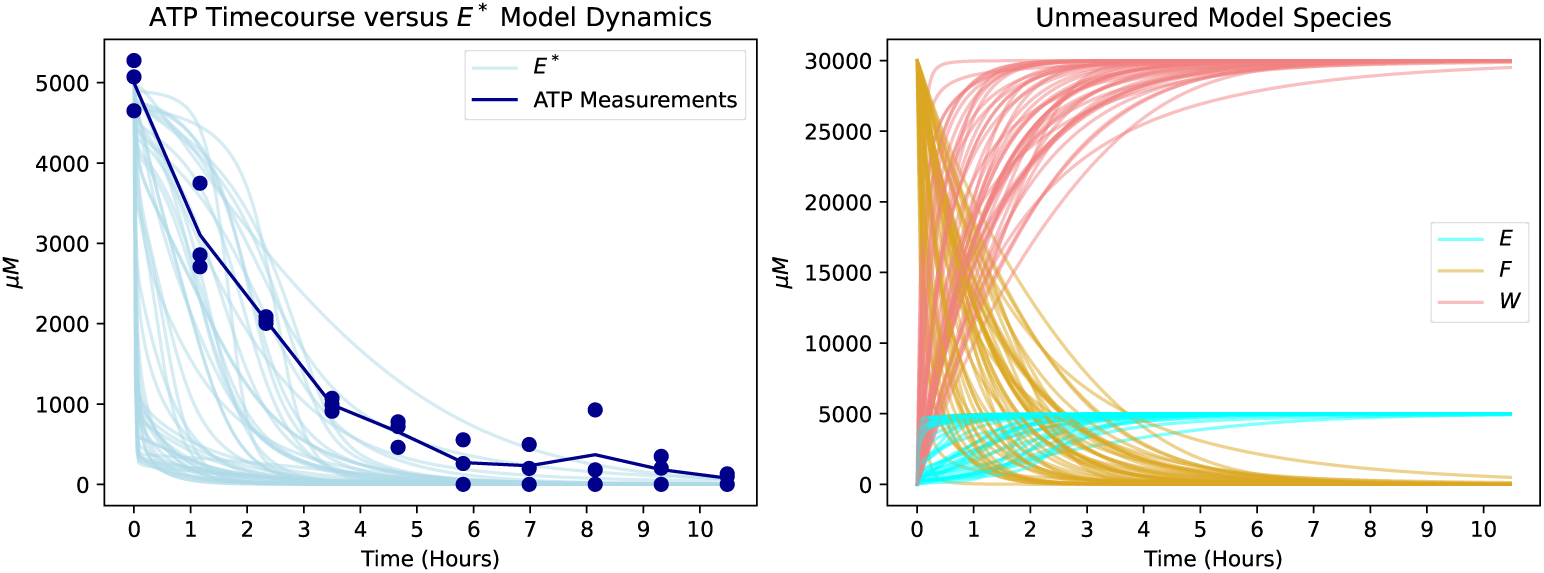
100 best parameter sets simulated from the baseline condition. Left: comparison between model simulations of the species *E*^∗^ (light lines) and ATP timecourse data (dark dots). Right: simulated dynamics of the other model species *E*, *F*, and *W*.

**Table 2:**
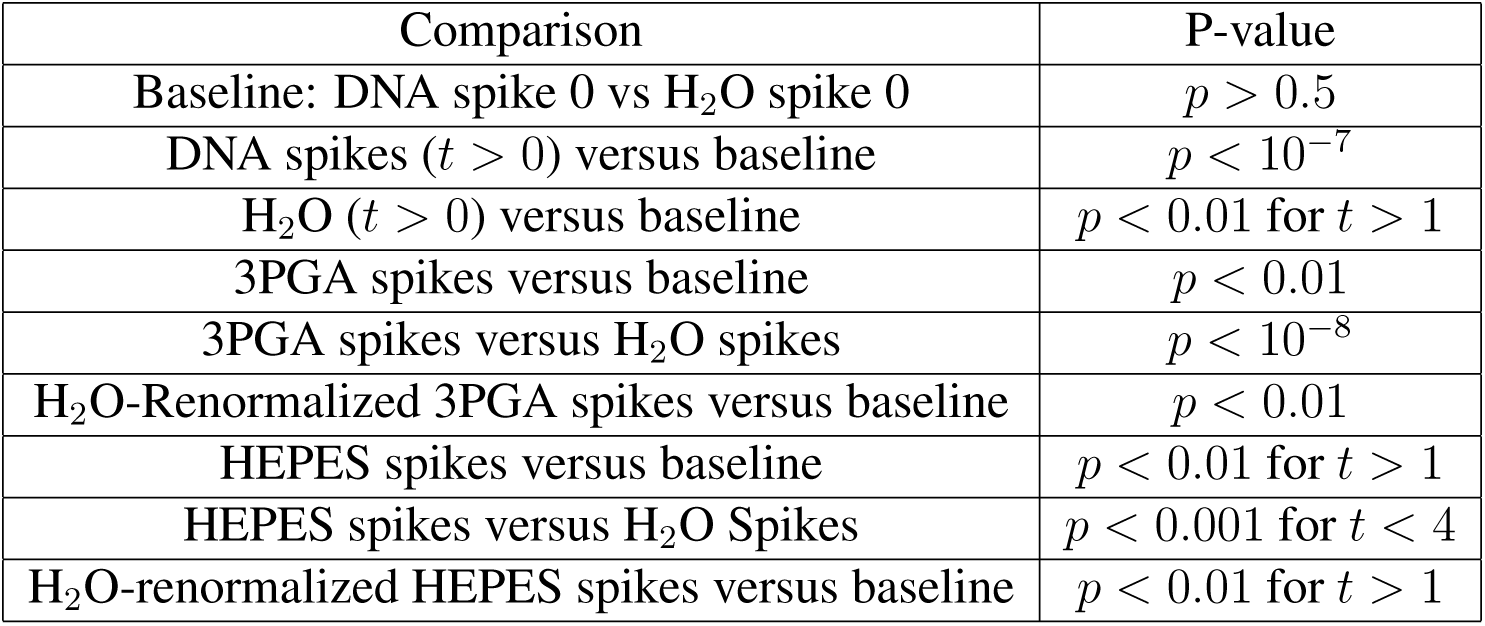
Statistics showing that the spiking experiments result in significantly different final protein expression compared to the baseline measurements or the H_2_O control spikes. H_2_O-renormalized measurements are scaled by the reduction in final output caused by H_2_O spikes in order to correct for changes in CFPS due to crowding effects. In some cases, only early *t < n* or later *t > n* spikes were found to be significant; if no spike number *t* is quoted, all spikes are significant.

Although the metabolomics measurements produced a significant amount of data, they are not sufficient to fully parameterize a systems-level model of *E. coli* lysate metabolism. Relatively recent *E. coli* flux balance analysis (FBA) models typically have on the order of over 2000 reactions between over 1000 metabolites involving over 1000 genes [35]. We expect that a systems-level lysate model would have to be of a similar size considering that most metabolic pathways appear to be active. Furthermore, the steady state assumption used in flux balance models is not valid for lysates. Therefore, we cannot ignore reaction rate parameters in favor of stoichiometric constraints and optimization used in FBA. Considering that we measured less than 30% of the metabolites likely active in cell lysate metabolism, we chose to take a phenomenological approach to modeling cell lysate metabolism [36, 37]. First, we note that cell lysate is powered by one dominate fuel source (in our experiments 3PGA). This fuel source is fed directly into the citric acid cycle and presumably used for mixed acid fermentation [35]. This allows cell lysate to regenerate ATP, GTP, NADH, and other energy-carrying molecules while simultaneously producing a whole host of other compounds and waste products. Additionally, we know that cell lysate energy buffer is fairly well optimized and that the addition of extra fuel, either initially or later in time, does not improve cell lysate performance and can even be toxic. Finally, if DNA is added to cell lysate after it has been incubated for a few hours, that DNA does not express. Jointly, these observations suggest that waste products build up in cell lysate, which shut down parts of its metabolism. This is a consistent with a previous finding that metabolic waste in CFPS systems is partially 3PGA-derived when 3PGA is used to regenerate energy [25].

To build a phenomenological model of CFPS, we require that our model matches the following experimental observations:

1. CFPS systems lose the ability to express DNA after 4-6 hours.
2. Adding additional fuel to CFPS systems either initially or after a delay does not improve CFPS performance.
3. Protein synthesis does not significantly impact cell lysate metabolism.

We build a simplified metabolic model consisting of four species: fuel *F*, energy carriers *E*^∗^ (activated, e.g. ATP), *E* (depleted, eg. ADP), and waste *W*, which interact via the following reactions using Hill function kinetics reminiscent of Michaelis-Menten enzyme kinetics [38]:

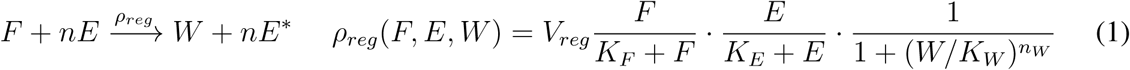

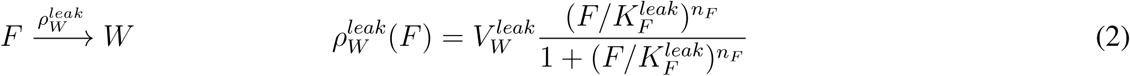

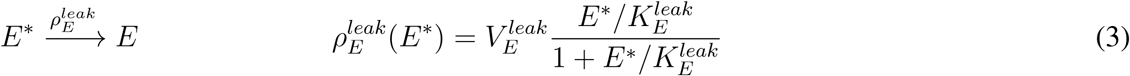

The first reaction (1) represents energy carrier regeneration from fuel. Here, *n* is a constant that depends on the particular fuel source (we used *n* = 3 because 1 molecule of 3PGA produces 3 ATP from 3 ADP but recognize that this constant is somewhat arbitrary for a non-mechanistic model). The rate function is given by a product of Hill function with a maximum velocity *V_reg_*. The first term represents the requirement that *F* be present for regeneration to occur. The second term represents the requirement that *E* be present for regeneration to occur. The third term represents the negative feedback due to build up of waste *W* which eventually shuts off metabolism.

The second reaction (2) represents the degradation of fuel *F* into waste products *W* without regeneration of any energy carrier *E*^∗^. Because the strength of the negative feedback of *W* on the rest of the system can be tuned via the constants *n_W_*and *K_W_*, this reaction effectively includes all utilization of the fuel for non-regenerative purposes, only some of which will produce toxic waste. The constants 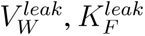 and *n_F_* are the maximum velocity of this reaction, the fuel concentration where this reaction reaches half its maximum velocity, and a Hill coefficient representing how sharply this process turns on as additional fuel is added to the system. Intuitively, we expect this reaction to model the fact that adding additional fuel to the CFPS system simply results in the accumulation of additional waste without improving regeneration.

The third reaction (3) represents the innate use of energy carriers *E*^∗^ by the cell lysate. The parameters 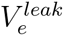 and 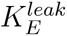 denote the maximum velocity of this reaction and how it depends on the amount of *E*^∗^ available. Intuitively, we expect this reaction to model the majority of energy utilization in CFPS.

These three reactions can then be combined with a CFPS module representing gene expression. For the purposes of this work, we coupled the metabolism model to a one-step gene expression model representing constitutive deGFP production:

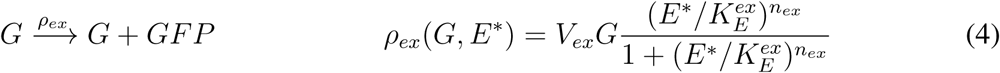

Here, *V_ex_* is the maximum rate of gene expression per unit of *G*. The Hill function term with parameters 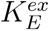 and *n_ex_* describe how gene expression depends on the available energy carriers *E*^∗^. We note that saturation of gene expression and explicitly modeling transcription and translation could easily be included in this framework, but would require additional data to determine the necessary parameters [39]. Additionally, in a more complex model, transcription and translation could consume energy. For example, the methods used in [40] would likely work in conjunction with this metabolism model provided data were collected to parameterize the entire system.

We fit the phenomenological model to the spiking time-course data using Bayesian parameter inference (see Methods). This resulted in a moderately robust fit of our model to the DNA spiking experiments and the baseline measurements. However, the 3PGA spiking experiments were not well fit by the model. We suspect this is because the effect size of 3PGA spikes is small resulting in 10% or less reduction on CFPS. We believe that this fit could be improved with more measurements involving larger titrations of 3PGA. A comparison of the mean spiking data with simulations from the top 100 best sets of model parameters can be seen in Figure 2.

### Model can be adapted and fine-tuned to fit and predict new experimental data

As our model fit the data from our baseline and DNA spiking experiments reasonably well, we were next interested in exploring the generalizability of our model. In particular, we were encouraged by the recent successes of foundation models in biology, where a base model trained on large amounts of data can be further “fine-tuned” to make it more predictive for a specific application, by retraining the model on a more relevant data set [41–44]. In our case, we wondered whether our model could be readily retrained to fit or even predict protein expression dynamics in new CFPS contexts, including new batches of cell lysate, different experimental initial conditions, and in the presence of complex biochemical interactions.

We first sought to determine whether the model architecture of our chemical reaction network was sufficiently expressive to fit a broad range of protein expression dynamics. Using a previously published data set, we first explored whether our model could be adapted to fit the new experimental data, which stems from CFPS systems using various fuel types and concentrations for energy regeneration, different concentrations of Mg^2+^, multiple cell lysate batches, different promoters for deGFP expression, and different DNA concentrations [25]. To perform the fine-tuning, we attempted to fit 576 individual time trajectories of deGFP expression by fine-tuning (i.e., re-inferring) 4 of the 13 total model parameters and fixing the remaining 9 parameters’ values to the median value of their posterior distributions (Figure S1A). These parameters – 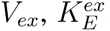, *K_W_*, and *n_W_* – were chosen because they were suspected to most likely be affected by different fuel sources, cell lysate batches, and promoters. The parameters *V_ex_* and 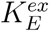 represent the maximum rate of protein production and the energetic requirement of producing the protein, respectively. *K_W_*and *n_W_*are parameters that determine the extent to which a given fuel’s waste negatively affects the rate of energy regeneration.

We found that the fine-tuning approach was very successful; the fine-tuned models better minimized error between model simulations and experimental data, compared to the base model where all parameters were used without further fine-tuning and only the initial conditions of the model were changed to reflect those of the experimental data (Figure S1B, S2). More importantly, this simple fine-tuning demonstrated that the metabolism model was sufficient to capture a broad range of endpoint deGFP concentrations and dynamics (Figure S1C-E), from deGFP expression that rapidly ceased at 5 hours to sustained deGFP expression that continued for 18 hours. Equally importantly, we found that these models could often be fine-tuned in as few as 500 steps and no more than 20000 steps, significantly fewer than the 100000 steps performed during the Bayesian parameter inference of the original model.

We next explored whether the fine-tuning approach could be extended to a model that was slightly different from the one it was originally trained on. In the previously published data we used [25], we noticed a DNA saturation effect in CFPS systems, such that after a certain concentration of DNA, deGFP endpoint yields would not change with increasing DNA concentration. Furthermore, in some systems, additional DNA resulted in reduced deGFP yields, perhaps due to a resource competition between transcription and translation. Both of these phenomena seemed to depend heavily on the particular Mg^2+^ and 3PGA concentrations used for the set of experiments. Our original CFPS model did not account for this effect, so we wondered if the fine-tuning approach could be used as before, albeit with (1) a new reaction propensity for deGFP expression and (2) when trying to fit multiple time trajectories instead of a single time trajectory (Figure 4A). For this new fine-tuning workflow, we re-inferred the same 4 parameters and newly inferred 4 more parameters resulting from the new reaction propensity. We also provided 5 time trajectories of deGFP expression corresponding to 5 different DNA concentrations for each of 36 different fine-tuning cases resulting from different combinations of Mg^2+^ and 3PGA concentrations.

**Figure 4:**
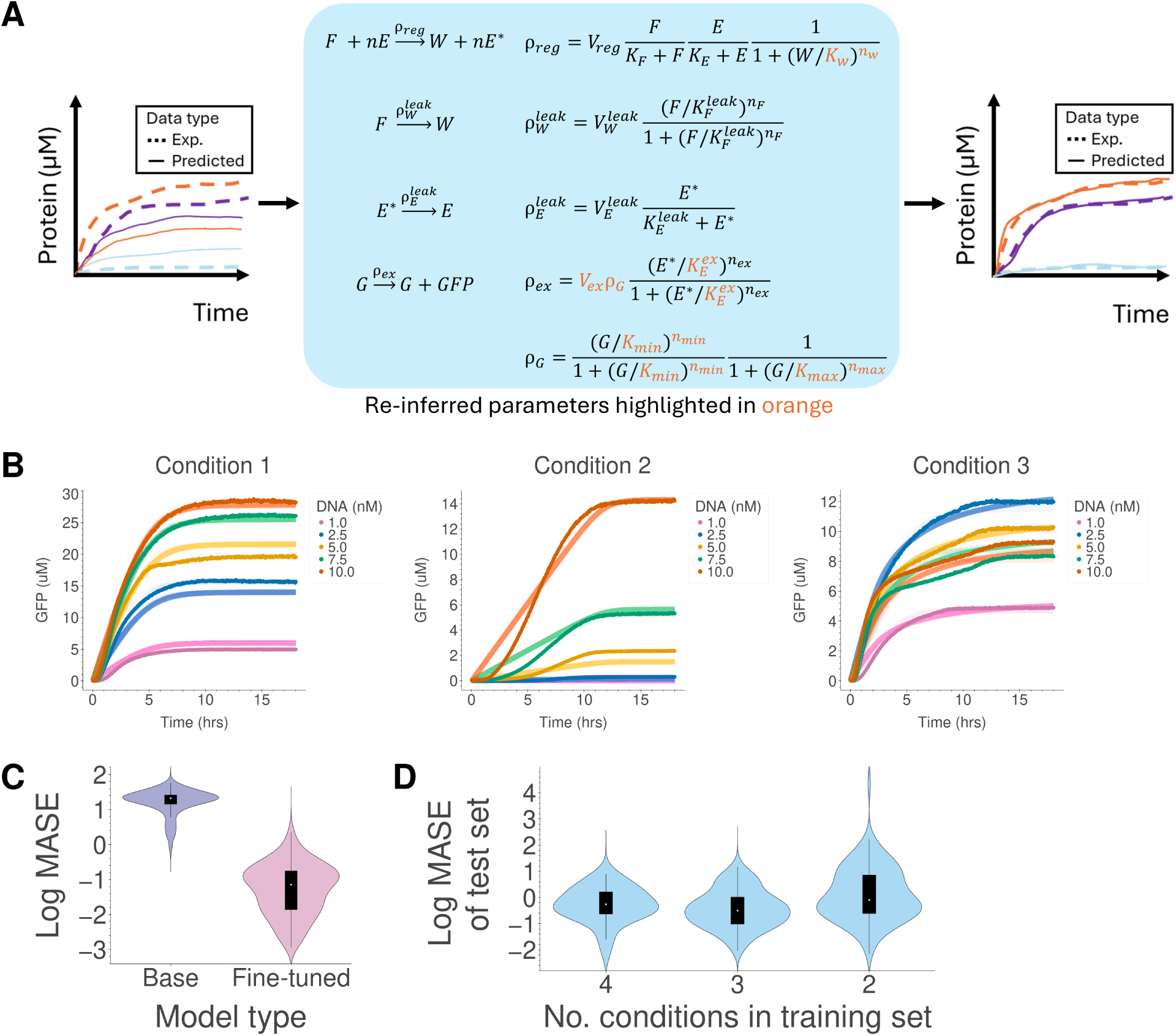
Fine-tuning a model with a modified reaction propensity. (a) Overview of computational workflow. Four parameters (highlighted in orange) of the already fitted base model were re-inferred and 4 parameters were newly inferred to better fit the model to new experimental data. Multiple time trajectories corresponding to various [DNA] for a given set of [3PGA] and [Mg^2+^] were provided for the fine-tuning. (b) Comparison of experimental data (dark points, average of 3 replicates) and model predictions (solid lines) after fine-tuning for 3 different experimental conditions: 5 mM 3PGA and 0 mM Mg^2+^ (Condition 1), 45 mM 3PGA and 0 mM Mg^2+^ (Condition 2), and 20 mM 3PGA and 6 mM Mg^2+^ (Condition 3). (c) Violin plots of log MASE values (see Methods) before and after fine-tuning the model on new data. Each violin represents a distribution of 180 MASE values (5 [DNA] x 6 [3PGA] x 6 [Mg^2+^]). For each of 36 fine-tuning cases, all 5 time trajectories’ data (corresponding to DNA concentrations of 1 nM, 2.5 nM, 5 nM, 7.5 nM, and 10 nM for a particular set of [3PGA] and [Mg^2+^]) were provided as both training data and test data. (d) The violins correspond to MASE values where the training and test data were split. For each violin, data corresponding to the following [DNA] was used for the test set (and withheld in the training set) for each set of 3PGA and Mg^2+^ concentrations: 5 nM DNA (left violin); 2.5 nM DNA and 7.5 nM DNA (middle violin); and 2.5 nM, 5 nM, and 7.5 nM DNA (right violin).

As before, we found that the fine-tuning approach was successful; the fine-tuned models better minimized error between model simulations and experimental data compared to the base model (Figure 4B,C). The fine-tuned models were successful at fitting a broad range of deGFP concentrations and dynamics (Figure 4B, S4) as well as capturing nonintuitive trends, such as deGFP expression being maximized at a DNA concentration that was not the highest used for that experiment (Figure 4B, Condition 3). The experimental conditions where the fine-tuning did not work well corresponded to experiments with initial conditions of low 3PGA and high Mg^2+^ concentrations, which resulted in biphasic protein expression that the model was not capable of capturing (Figure S4). Even in the presence of unusual dynamics, however, the model often succeeded at matching the endpoint deGFP yields obtained for different initial concentrations of DNA (Figure S4).

To determine whether the model could not only fit data but also predict unseen data, we next explored fine-tuning where the same experimental data was split into training and test sets. Here, the fine-tuning was performed on training data only, after which the test data was compared to model predictions for those experimental conditions. Due to the reaction propensity of the protein expression reaction, however, we could not simply randomize the assignment of data into training and test sets. As mentioned above, deGFP yields first increased, then decreased, with increasing DNA concentration in a CFPS system, with the relationship between endpoint deGFP yields and initial DNA concentration resembling the shape of a bandpass filter. Thus, providing concentrations on the end of the range of concentrations used (i.e., 1 nM and 10 nM DNA) was necessary to ensure the shape of the function could be maintained. So, we split the data into training and test sets in the following manner. For each fine-tuning case, we first reserved the time-course deGFP dynamics corresponding to 5 nM DNA for the test set, and used the remaining 4 DNA concentrations’ data (1 nM, 2.5 nM, 7.5 nM, and 10 nM) for the training data used during fine-tuning. We next repeated this procedure, albeit using data corresponding to 2.5 nM and 7.5 nM DNA for the test set, and using the remaining 3 DNA concentratons’ data (1 nM, 5 nM, and 10 nM DNA) for the training set. Finally, we repeated this procedure again, albeit using the data corresponding to 2.5 nM, 5 nM, and 7.5 nM DNA for the test set, and using the remaining 2 DNA concentratons’ data (1 nM and 10 nM DNA) for the training set. The error between model prediction and experimental data for the test sets are shown in Figure 4D (see Figure S3 for comparison between Figure 4C and Figure 4D).

We found that fine-tuned models were generally successfully at capturing the shapes of the deGFP time-course dynamics, although they often struggled to recapitulate the exact concentrations achieved experimentally (Figure S5-S7, compared to Figure S4). This issue was exacerbated, as expected, when less data was provided in the training set. It is worth noting that the fine-tuned models were surprisingly successful at qualitatively predicting some phenomena, including no deGFP expression in a TX-TL system with 2.5 nM DNA (Figure S6, at 45 mM 3PGA and 0 mM Mg^2+^), relatively high deGFP expression with a different TX-TL system with 2.5 nM DNA compared to other DNA concentrations (Figure S6, at 5 mM 3PGA and 2 mM Mg^2+^), and comparable deGFP expression in systems with 7.5 nM and 10 nM DNA (Figure S7, at 45 mM 3PGA and 10 mM Mg^2+^).

### Adapted and fine-tuned model can be used to gain novel experimental insights

Having confirmed that the fine-tuning approach was successful at fitting and predicting deGFP protein expression dynamics, we next wondered whether the fine-tuning approach could be used to gain experimental insights in a novel experimental system. The study whose data we were using for fine-tuning had found a compensatory interaction between Mg^2+^ and 3PGA in CFPS systems, where too much of either component reduced protein yields, but enough of both was sufficient to revive protein expression [25]. We wondered if our model could be adapted and our fine-tuning approach applied to qualitatively model this phenomenon and explore potential causes for this phenomenon. For this fine-tuning workflow, we first added Mg^2+^ as a new species to the model, modified our protein expression reaction to account for the inhibitory effects of Mg^2+^ on deGFP expression, and added a reversible reaction for binding of Mg^2+^ to waste in the model (Figure 5A). Next, we re-inferred the same 4 parameters as in Figure 4 and newly inferred 4 more parameters resulting from the new reaction propensity and new reactions added to the model (Figure 5A). Unlike the previous fine-tuning case, however, where we fine-tuned separately for small sets of experimental data, for each of two fine-tuning cases, we provided 36 time trajectories of deGFP expression corresponding to 6 different 3PGA concentrations and 6 different Mg^2+^. We found that while our fine-tuning approach did not do well quantitatively, it captured the experimentally observed deGFP dynamics qualitatively at low 3PGA concentrations (Figures S8, S9). More interestingly, the model captured the same general endpoint trends in deGFP yields as observed experimentally (Figure 5B,C) in the previously published study [25].

**Figure 5:**
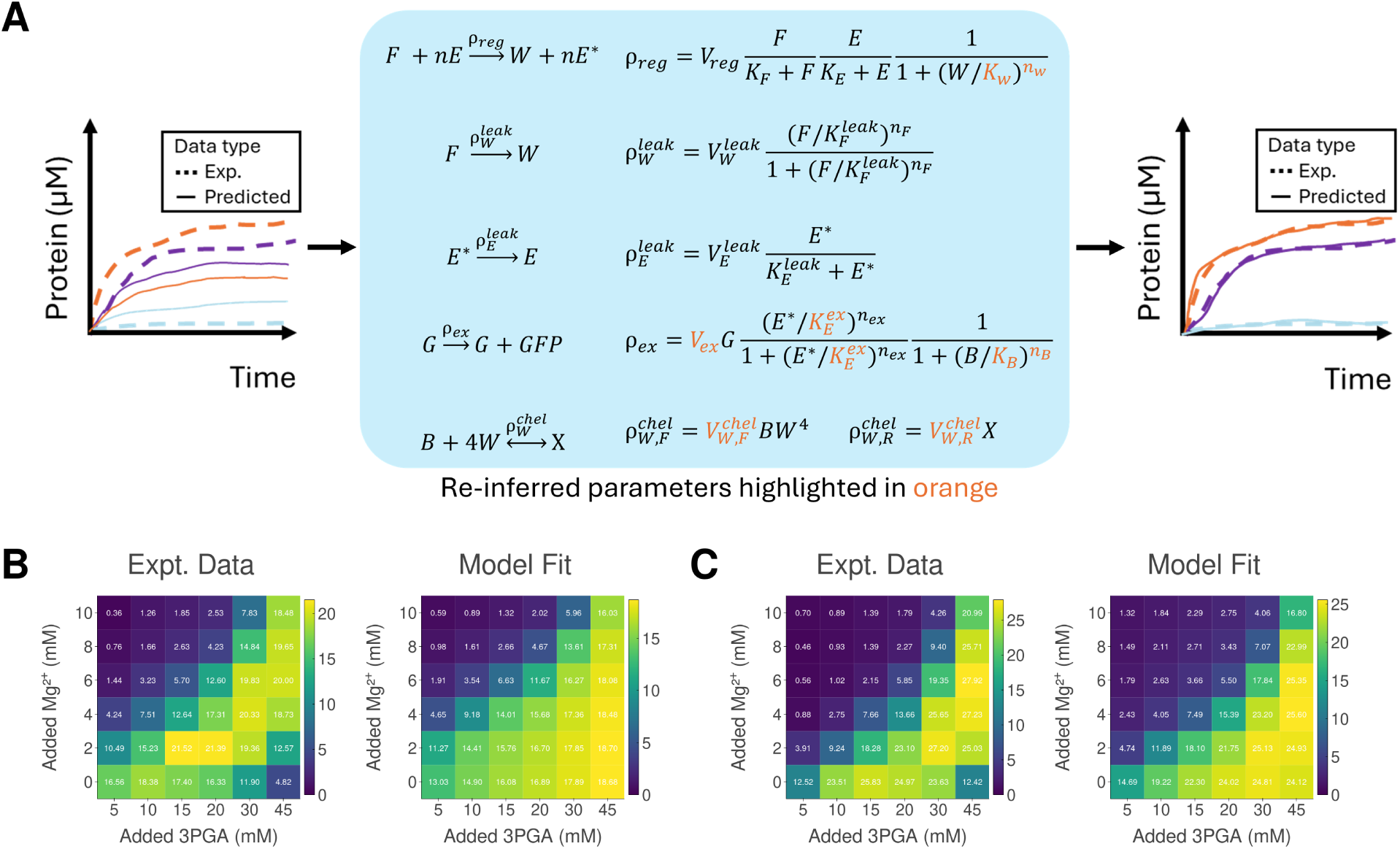
Fine-tuning a model with an additional reaction. (a) Overview of computational workflow. Four parameters of the already fitted base model – *V_ex_*, 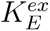, *K_W_*, and *n_W_* – were re-inferred and four parameters – *K_B_*, *n_B_*, 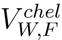 and 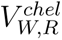 – were newly inferred to better fit the model to new experimental data. 36 deGFP time trajectories corresponding to 36 sets of 3PGA and Mg^2+^ concentrations were provided for the fine-tuning for each fine-tuning case. (b) and (c) show endpoint deGFP comparisons between experimental data and model fit after fine-tuning for Lysate Batches 1 and 2 from the experimental data previously published [25].

By modeling species not measured experimentally, we were also able to glean some new experimental insights. First, we noticed that the two batches of cell lysate, whose deGFP dynamics were used to parameterize the model separately during their respective fine-tunings, metabolized fuel at different rates, with Lysate Batch 2 metabolizing fuel slower and generating waste slower than Lysate Batch 1 (Figures S10, S11). Meanwhile, however, energy concentrations were consistent across the two batches of lysate. This seemed to suggest that the higher deGFP yields observed in Lysate Batch 2 were due to lower waste generation rather than improved energy regeneration, which is consistent with our original hypothesis that waste generation limits protein expression to a greater extent than energy regeneration. Second, our model simulations also suggested the Mg^2+^-W binding reaction had a fast reaction rate (Figures S10, S11). This is consistent with a previous paper that found that Mg^2+^ had to be repeatedly titrated into a CFPS reaction, rather than added in a high amount all at once, to improve protein expression yields, since adding too much Mg^2+^ at once inhibited protein expression [45]. Finally, the model simulations suggested that adding additional fuel above 15-20 mM 3PGA to a CFPS reaction does not appreciably increase the concentration of energy available in the system (Figures S10, S11). This is consistent with our own data (Figure 2), where we also found that the addition of 3PGA to a CFPS system with 30 mM 3PGA did not improve deGFP yields, and with a previous paper that suggested that most of the energy regenerated during CFPS is diverted towards metabolic pathways competing with CFPS for energy [25].

## Discussion

The metabolomics measurements in this work present strong evidence of the complexity underlying cell lysate-based CFPS metabolism. In particular, we found that a large abundance of initial fuels powers metabolic activity across the majority of *E. coli* metabolic pathways. We hypothesize that this metabolism results in the accumulation of multiple waste products which ultimately inhibit CFPS. Identifying which metabolic end-products are responsible for this inhibition and how metabolic activity may vary with different CFPS conditions remains a challenge. However, based upon our metabolic data, we make several observations about the activity of specific pathways in cell lysate metabolism and concrete hypotheses about potential waste sources.

### Fuel consumption produces phosphate

As expected, 3PGA (3-phospho-D-glycerate) is consumed along with cellular PEP in order to generate ATP. ATP is consumed by reactions required for protein synthesis (e.g., tRNA charging) which generates products such as AMP. This process also produces inorganic phosphate, which has previously been reported as a potential waste product responsible for diminishing CFPS [45, 46].

### Magnesium ions

The compensatory interaction between Mg^2+^ and 3PGA in previously published data [25] was unexpected and suggests that CFPS systems operate optimally within a narrow range of Mg^2+^ concentrations. On the higher end of Mg^2+^ concentrations, the sensitivity of cell lysate to this ion may be due to phosphoglycerate kinase. This enzyme is inhibited by high Mg^2+^ concentrations. When inhibited, this would prevent the regeneration of ATP regeneration via glycolysis. Meanwhile, on the lower end of Mg^2+^ concentrations, Mg^2+^ is rapidly chelated by waste products and cannot perform vital tasks for CFPS, including serving as a cofactor for RNA polymerase and other enzymes constituting TX-TL machinery. The ability of Mg^2+^ to chelate a waste product also suggests that investigating ions that bind to Mg^2+^ in CFPS reactions (e.g., pyrophosphate) might point to additional waste product sources.

### NADH consumption

The majority of NADH is expected to be oxidized to NAD+. However, the NAD+ concentration remains constant, presumably via allosteric regulation of enzymatic activity. Several NAD+ pathways are potentially responsible for the breakdown of NAD+, ensuring its homeostatic concentration. This includes the NAD salvage pathway IV, which operates in reverse, generating 1-(*β*-D ribofuranosyl)nicotinamide.

### TCA cycle

We suggest that reactions catalyzed by isocitrate dehydrogenase and succinate:quinone oxidoreductase [(SdhA)(SdhB)(SdhC)(SdhD)]3 – required for a functional TCA cycle – are significantly down-regulated in cell-free metabolism:

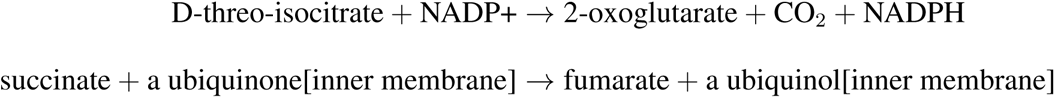

This down-regulation could lead to a build of intermediates (e.g., succinate) and a reduction in reducing power for ATP synthesis. From our data, we cannot precisely determine the reason why these reactions are inhibited; it could be due to missing or inactive enzymes, missing cofactors, or other reasons. One possibility is that succinate:quinone oxidoreductase [(SdhA)(SdhB)(SdhC)(SdhD)]3 is a complex membrane bound heterotetramer which might be inactivated and/or diluted during lysis and centrifugation. Another possibility is due to a low abundance of NADP. While we didn’t quantify NADP, it is not added exogenously to CFPS systems and might explain why the reaction catalysed by isocitrate dehydrogenase is inactive. Notably, we also could not quantify D-threo-isocitrate and therefore the proceeding reaction could be inactive, explaining the accumulation of cis-aconitate and depletion of 2-oxoglutarate.

### Glycine biosynthesis

We noticed that the concentration of L-glycine more than doubles during the CFPS reaction. Given the relatively high concentration of 3PGA, it is possible that this molecule is being utilized along with glutamate to synthesise glycine. This would divert energy away protein synthesis, representing an inefficiency in the system.

### Purine degradation

As previously noted, we observe an increase in AMP from reactions required for protein synthesis. We also observe a transient build-up of GMP during the reaction. There are several known reactions involving GMP, including reactions which consume ATP. It is feasible that these reactions operate in reverse to compensate for a reduction in ATP during the reaction. In the case of AMP and GMP we see further breakdown with a build-up of xanthine and ultimately ureate/uric acid. We note that uric acid is not processed by *E. coli* metabolism and therefore represents a final waste product which may inhibit CFPS. Uric acid along with NH^+^ ions generated during these processes could function to inhibit or denature enzymes.

Considering the great many uncertainties we have with regards to the interplay of CFPS and metabolism, as illustrated by the above discussion, we opted to simplify the complexity of cell lysate metabolism in order to arrive at a useful model for CFPS. To do this, we coarse-grained metabolism into three abstract species: fuel molecules, energy carriers, and waste molecules. We were then able to phenomenologically model the feedback these abstract metabolic species have on CFPS via a simple chemical reaction network which we fit to CFPS data. We then show that this mechanistically motivated model is expressive enough to capture the expression dynamics of hundreds of cell lysate conditions when a subset of its parameters are fine-tuned on condition-specific CFPS data.

Our approach demonstrates a novel modeling strategy where we trained a coarse-grained metabolic model on a large dataset and fine-tuned a subset of its parameters on specific lysate conditions. This methodology is similar to the approach taken in [47] where parameter non-identifiability is exploited in order to correct for batch effects in cell lysates. Our approach is also inspired by modern machine learning methods; foundation models, both in biological applications and more broadly, typically consist of a deep learning model trained on large data sets which are then frequently finetuned on more specialized datasets in order to perform specific tasks [41–44]. In biology, these large data sets can include omics data (transcriptomics, genomics, metabolomics, etc.), language data (from the Internet and other sources), and even images. We have adapted this approach to use an interpretable mechanistic model fit to a carefully constructed panel of CFPS time-coarse data and then fine-tuned to model various CFPS conditions. As our models are simple and mechanistic, we can use our model to provide experimental insights in addition to predictability, a capability not typically possible with “black box” deep learning models. In the example shown in Figure 4, we show that model can be readily adapted to fit and predict protein expression dynamics previously unseen in the original training data. In the example shown in Figure 5, we show that this model can additionally be used to gain experimental insights into cell-free metabolism and guide future experimental efforts aimed at characterizing and improving cell-free metabolism to benefit CFPS. By providing detailed metabolomics insights into CFPS systems and computational frameworks that can be used to model protein expression dynamics in the presence of this metabolic complexity, this work provides experimental and computational tools for understanding the complex interplay between cell-free metabolism and CFPS. This work forms a solid step towards improving the predictability of CFPS systems, which will be necessary for future cell-free applications aimed at building complex systems, from multi-layered biological circuits to synthetic biological cells.

## Methods

### CFPS Reactions

#### Cell Lysate Preparation

Crude cell lysates were all prepared according to the protocol [2]. Briefly, all batches consist of 6 x 0.75L *E. coli* (BL21 Rosetta) cultures grown in 2xYT media supplemented with phosphate buffer to an OD of ∼ 3 seeded from 7.5mL of culture grown for 8 hours. Cells are then pelleted and rinsed twice with S30 buffer before being lysed either via sonication [48] or French press [49]. Post lysis, the cell lysate is purified via two rounds of centrifugation at 30000*g*. All batches were then incubated for 1 hour at 37^◦^C in a runoff reaction. Some batches were dialyzed using 10k MWCO Dialysis Cassettes in S30B buffer. Finally, the cell lysate was flash frozen in liquid nitrogen for later use.

#### Energy Buffers and Additional Additives

Frozen cell lysate samples were thawed and mixed with an energy buffer which uses 3PGA as the main fuel source and also supplements NTPs, amino acids, and a variety of co-factors listed in [49]. Cell lysate (33% by volume) and energy buffer (25% by volume) was mixed together with Mg-Glutamate (to a concentration of 5 or 10 mM), K-glutamate (100 mM for the Metabolomics measurements, 200 mM for spiking experiments), water, and, in some experiments, a constuitively expressing deGFP plasmid [50] MIDI prepped from an overnight cell culture and eluted in water.

### Metabolomics

#### Sample Preparation

The only element of the energy buffer which is varied across these experiments is magnesium glutamate which is known to strongly effect CFPS [49]. Some samples also received a constiuitively expressing deGFP plasmid at a final concentration of 4 nM. Cell free reactions (volume = 150 uL) were incubated at 37^◦^C for 0, 3, 6, or 12 hours in a sealed 96 well plate. Some samples received exposure to extra oxygen by periodic unsealing and resealing of the 96 well reaction plate during incubation. Reactions were stopped via the addition of cold methanol (volume = 37.5 uL) [51] and immediately flash frozen in liquid nitrogen. All the sample conditions are listed in Table 1.

#### Mass Spectrography

Frozen samples were shipped on dry ice to Metabolon for Ultrahigh Performance Liquid Chromatography-Tandem Mass Spectroscopy (UPLC-MS/MS). We quote their technical description of their pipeline: “All methods utilized a Waters ACQUITY ultra-performance liquid chromatography (UPLC) and a Thermo Scientific Q-Exactive high resolution/accurate mass spectrometer interfaced with a heated electrospray ionization (HESI-II) source and Orbitrap mass analyzer operated at 35,000 mass resolution. The sample lysate was dried then reconstituted in solvents compatible to each of the four methods. Each reconstitution solvent contained a series of standards at fixed concentrations to ensure injection and chromatographic consistency. One aliquot was analyzed using acidic positive ion conditions, chromatographically optimized for more hydrophilic compounds. In this method, the lysate was gradient eluted from a C18 column (Waters UPLC BEH C18-2.1x100 mm, 1.7 µm) using water and methanol, containing 0.05% perfluoropentanoic acid (PFPA) and 0.1% formic acid (FA). Another aliquot was also analyzed using acidic positive ion conditions, however it was chromatographically optimized for more hydropho-bic compounds. In this method, the lysate was gradient eluted from the same afore mentioned C18 column using methanol, acetonitrile, water, 0.05% PFPA and 0.01% FA and was operated at an overall higher organic content. Another aliquot was analyzed using basic negative ion optimized conditions using a separate dedicated C18 column. The basic lysates were gradient eluted from the column using methanol and water, however with 6.5 mM Ammonium Bicarbonate at pH 8. The fourth aliquot was analyzed via negative ionization following elution from a HILIC column (Waters UPLC BEH Amide 2.1x150 mm, 1.7 µm) using a gradient consisting of water and acetonitrile with 10 mM Ammonium Formate, pH 10.8. The MS analysis alternated between MS and data-dependent MSn scans using dynamic exclusion. The scan range varied slighted between methods but covered 70-1000 m/z.” Finally, Metabolon used their proprietary software and library of more than 3300 purified samples to identify different metabolites and normalize the resulting data. We emphasize that even with these methods the metabolite abundances reported are unitless and can only be quantified relatively.

### Spiking Experiments

Spiking experiments were conducted with 5 replicates on a 384 well plate using 10 µL reaction volumes. Cell lysate and buffer were mixed together and loaded by hand into each well. Then a Labcyte Echo liquid handling robot was used to load DNA and water to a volume of 9 uL. The following components were added as 1 µL spikes using the Labcyte Echo in different experiments: DNA (final concentration 2.4 nM); water; 3PGA (30 mM) and HEPES (50 mM). deGFP fluorescent data was collected from a Biotek plate reader using a gain of 61, excitation wavelength of 485 nm, and emission wavelength of 515 nm. Raw data can be seen in Figure 6. The DNA spikes used a different BioTek instrument than the other spikes, therefore data was normalized so that the two baseline measurements (shown in red in Figure 6) have the same steady final mean (these two conditions contain identical reagents). The other measurements have been calibrated using purified deGFP protein. Additionally, the HEPES and 3PGA spikes were normalized based upon the water spikes to correct for variation due to concentration changes. These different experiments have been statistically compared to each other in Table 2 using the Kruskal Wallis test to analyze variance in the final protein concentrations. Note that the optimal salt concentrations of 10 mM Mg and 200 mM K were chosen based upon a salt calibration screen for this particular cell lysate batch. The cell lysate used for these experiments was lysed with sonication and was not dialyzed.

**Figure 6:**
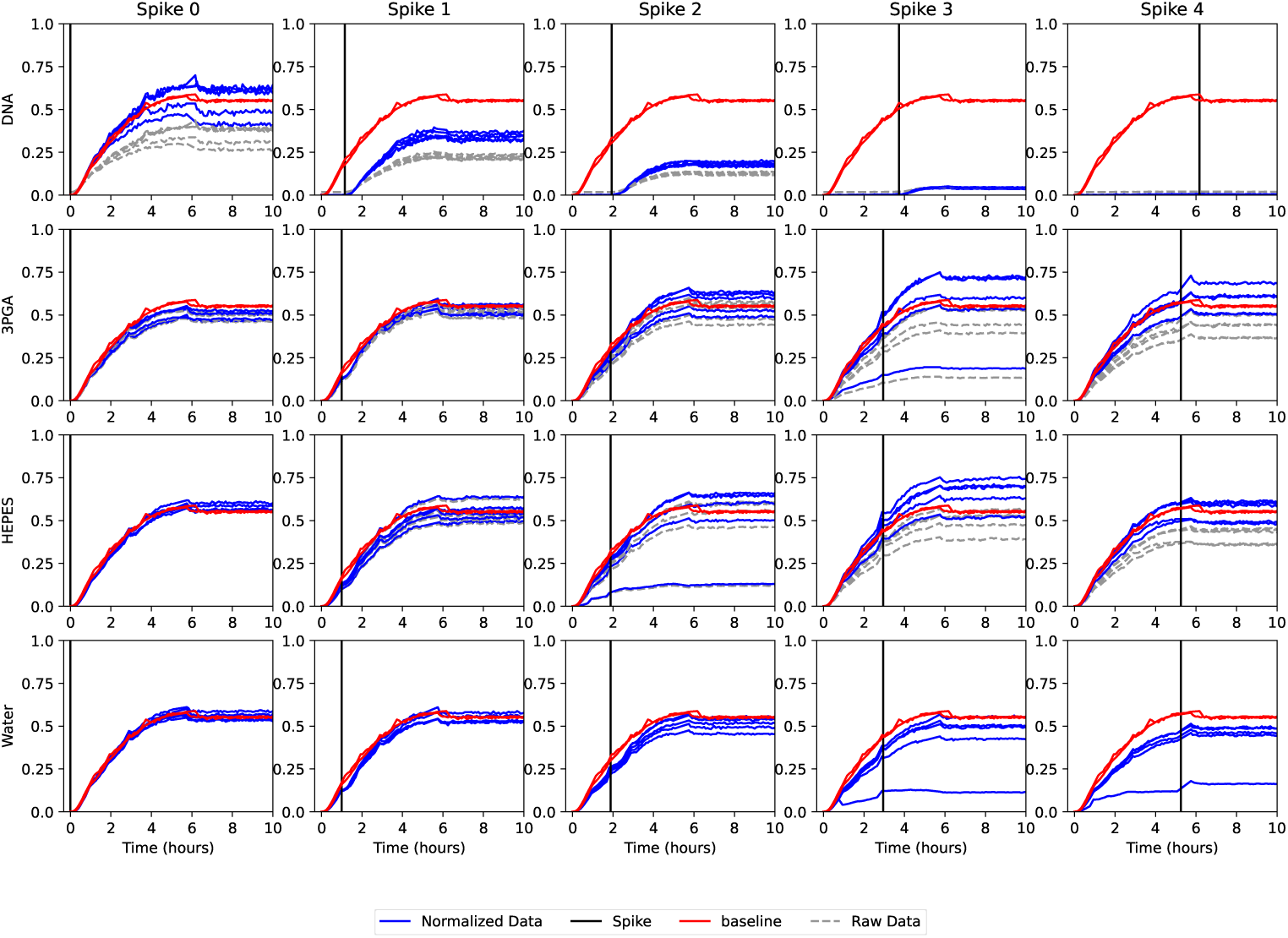
Spiking experiment raw data (gray) and renormalized data (blue). Red shows baseline measurements were computed from the DNA and Water Spike #0.

### ATP Time Course Measurements

ATP time course data was collected using the same cell lysate batch and buffer conditions as the spiking experiments. A 384 well plate was loaded with cell lysate and energy buffer and water (30 µL total reaction volume per well) and a liquid handling robot (Hamilton) was used to automatically measure the ATP levels using the Cayman Chemical Luciferase based ATP assay every hour for 10 hours (3 replicates per time point).

### Statistical Methodology

Statistics comparing measurements between individual metabolites were generated from the Kruskal-Wallis test (a non-parametric ANOVA) [52] between two or more sets of the same metabolite *m* in difference conditions. Specifically, the time course variation p-value for an individual metabolite *m* was generated from the Kruskal-Wallis test between four sets of measurements: one for each time point (pooled across all samples). The preparation method p-value for an individual metabolite *m* was generated from the Kruskal-Wallis test between different sets of preparation methods and/or cell lysate batches (pooled across all timepoints).

Other p-values reported were generated by combining p-values using Empirical Brown’s Method [32], an extension of Fisher’s method which takes into account covariances in the data used to generate the p-values. Specifically, time course variation p-values between different conditions (e.g. DNA vs No DNA) were generated by computing a p-value with the Kruskal-Wallis test for a metabolite *m* and timepoint *t* between the two conditions. The p-values for this metabolite and conditions across all timepoints were then combined with Empirical Brown’s Method to get a single time-varying p-value for that condition and metabolite.

Pathway level statistics were computed by combining either time course variation or prepara-tion p-values of many metabolites grouped together based on the pathway annotations provided by Metabolon using Empirical Brown’s Method. This methodology significantly enhances the statistical power of the data by allowing less significant variability to contribute to grouped meta-statistical p-values.

Significance testing was conducted at a threshold *p <* 0.01 rescaled for multiple testing using the Holm–Bonferroni correction. A total of 3280 metabolite-specific p-values were computed comparing different conditions of which 171 were found to be significant. Similarly, 550 pathway p-values were computed across different conditions of which 157 were found to be significant.

### Simulations and Parameter Inference

Models were produced using BioCRNpyler [53] saved as SBML files [54] and then loaded into Bioscrape [55] which uses the Emcee package [56] for black-box Bayesian parameter inference. The cost function used was the *L*_2_ norm between the mean simulated data for baseline measurements, DNA spike measurements, and 3PGA Spike measurements, specifically:

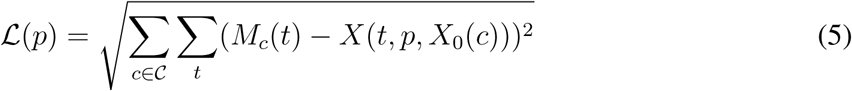

Here, L(*p*) is the likelihood of the parameters *p*. C is a set of all experimental conditions, and *M_c_*(*t*) denotes the measurement of condition *c* at time *t*. *X*(*t, p, X*_0_(*c*)) denotes a simulated trajectory of the CRN evaluated at time *t* using parameter *p* with initial condition *X*_0_(*c*) dependent on the specific measurement. Spiking was modeled by having time dependent rate constants which turn on at the spike times allowing a DNA or Fuel to flow into the system. Inference was split into two runs, the first with 50 walkers for 5000 steps and the second with 100 walkers for 10000 steps. Each step corresponds to simulating each walker with a given set of parameters at each different initial condition. This resulted in over ten million individual simulations with a total of 1,250,250 parameter combinations sampled. The likelihood of different parameter combinations can be visualized by looking at the marginals and pairwise marginals of the 13 model parameters as shown in Figure 7.

**Figure 7:**
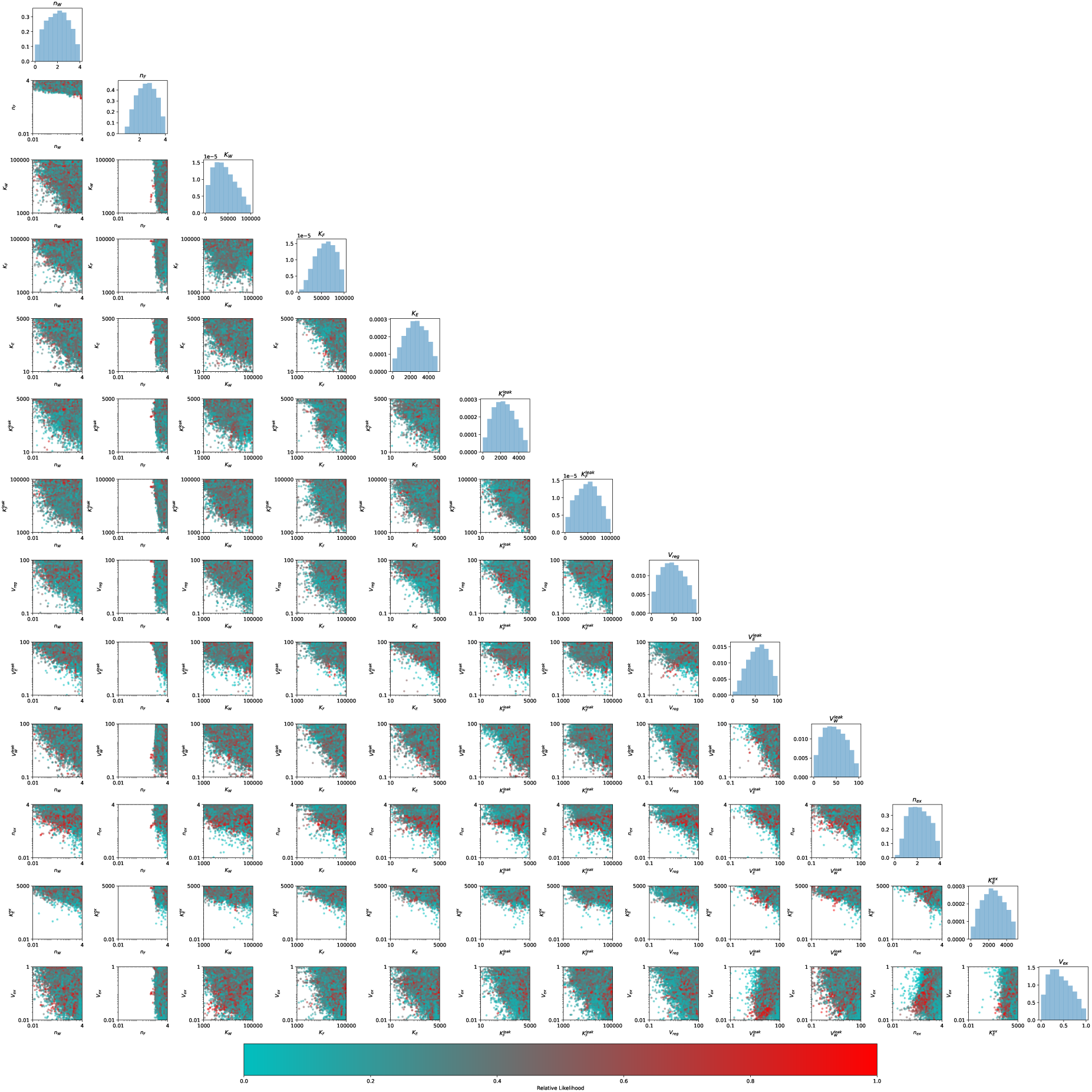
Parameter Posterior Visualization. Off-diagonal plots show the pairwise marginals of sampled parameters with the likelihood indicated by the color bar. Diagonal plots show the marginal distributions of each individual parameters.

### Model Recalibration

Model recalibration, or fine-tuning, was performed by first creating a model parameterized by the mean values of parameters of the top 100 parameter sets (as measured by parameter sets that resulted in the lowest error between model simulation and experimental data) from the posterior distribution of the Bayesian inference performed on the data shown in Figure 2. Next, any modifications to model propensities or new reactions were added to the model where applicable, with parameters for those additions hand-tuned to give the model a good starting point. Finally, the indicated parameters were then inferred/re-inferred by performing Bayesian inference with 100 walkers and a variable number of steps: 5000 steps for the fine-tuning described in Figure S1, 50000 steps for the fine-tuning described in Figure 4, and 50000 steps for the fine-tuning described in Figure 5. As before, models were produced using BioCRNpyler [53], saved as SBML files [54], and then loaded into Bioscrape [55] for inference.

### Mean Average Scaled Error (MASE) Plots

For figures where a MASE value was computed, the following approach was used. First, for each unique experimental condition, the model was simulated using the same initial conditions of model species (fuel (*F*), DNA (*G*), and NTP (*E*) concentrations) as those used in the experiment. Next, the percentage error between experimental data (averaged over 3 replicates) and model prediction was computed at each timepoint. Next, the mean value of these errors was computed to create a single MASE value. Finally, the distribution of MASE values corresponding to a given set of experimental conditions/data was displayed using a violin plot.

## Acknowledgments

We note that this work builds on preliminary results from W. Poole’s thesis [57]. We also note that the following authors completed this work at institutions different than the ones with which they are currently affiliated: Ankita Roychoudhury and William Poole contributed to this work while at the California Institute of Technology, and Matthew Haines contributed to this work while at Imperial College London. Research was supported by the following funding sources: the Army Research Office, accomplished under Cooperative Agreement Number W911NF-22-2-0210; the Institute for Collaborative Biotechnologies through contract W911NF-19-D-0001 from the U.S. Army Research Office; and the National Science Foundation award number 2152267. The views and conclusions contained in this document are those of the authors and should not be interpreted as representing the official policies, either expressed or implied, of the Army Research Office or the U.S. Government. The U.S. Government is authorized to reproduce and distribute reprints for Government purposes notwithstanding any copyright notation herein.

## Supporting information

**Figure S1:**
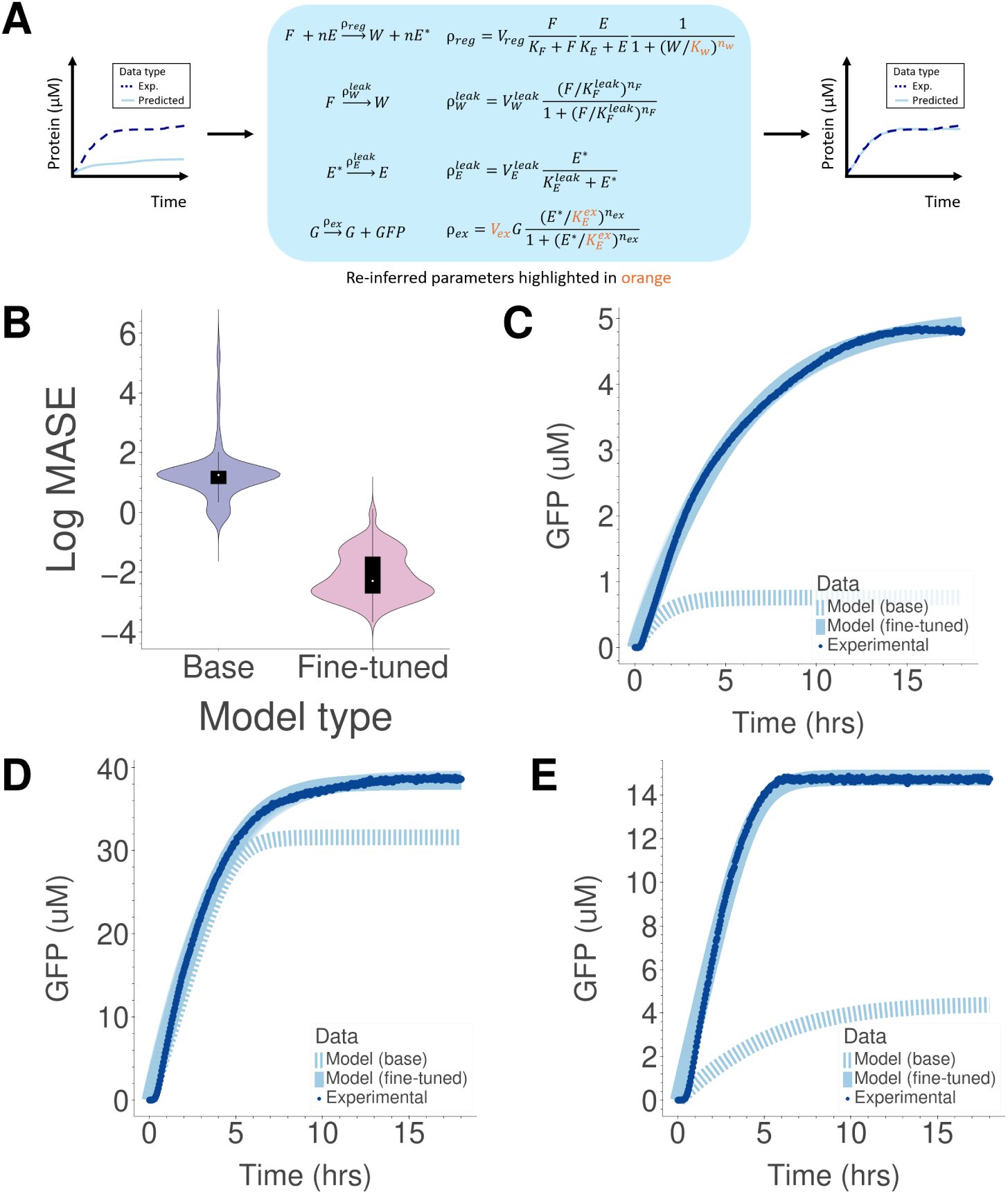
Verification of fine-tuning approach for individual experimental conditions. (a) Overview of computational workflow. Four parameters (highlighted in orange) of the already fitted base model were re-inferred to better fit the model to new experimental data. (b) Violin plots of log MASE values (see Methods) before and after fine-tuning the model (aggregated data from the plots shown in Figure S2). (c)-(e) show experimental data and model predictions before and after fine-tuning for three different experimental conditions: (c) Lysate 1, 30 mM maltose, 4 mM Mg^2+^, 5 nM DNA with P_OR1OR2_ promoter; (d) Lysate 2, 30 mM 3PGA, 4 mM Mg^2+^, 10 nM DNA with P_OR1OR2_ promoter; (e) Lysate 3, 20 mM 3PGA, 0 mM Mg^2+^, 5 nM DNA with P_T7_ promoter.

**Figure S2:**
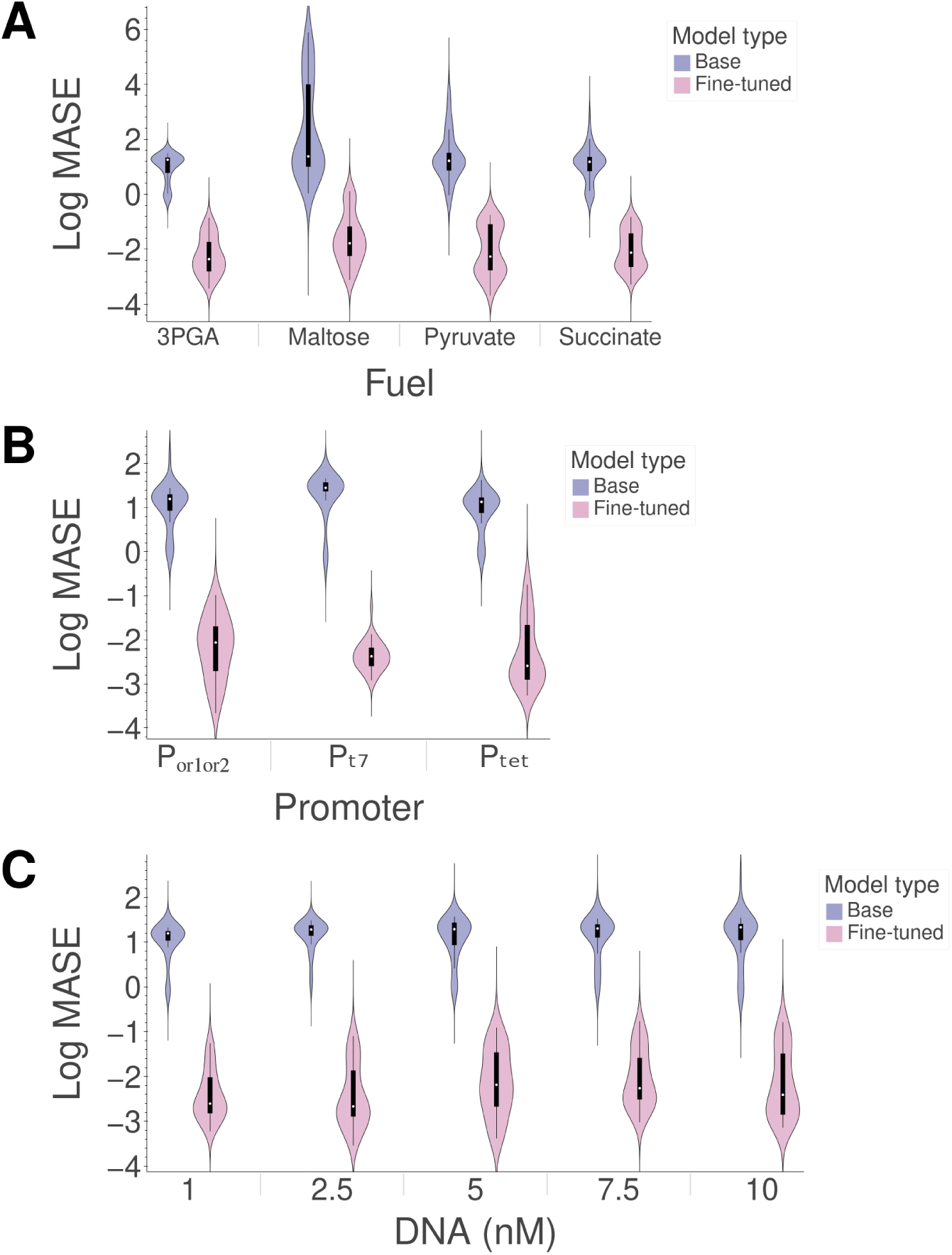
Violin plots of log MASE values before and after fine-tuning model. Each of the plots (a), (b), and (c) show data from different experiments. Each violin represents the distribution of the following number of unique experimental conditions: (a) 72 (2 lysate batches x 6 fuel concentration x 6 Mg^2+^ concentrations), (b) 36 (6 fuel concentration x 6 Mg^2+^ concentrations), and (c) 36 (6 fuel concentration x 6 Mg^2+^ concentrations).

**Figure S3:**
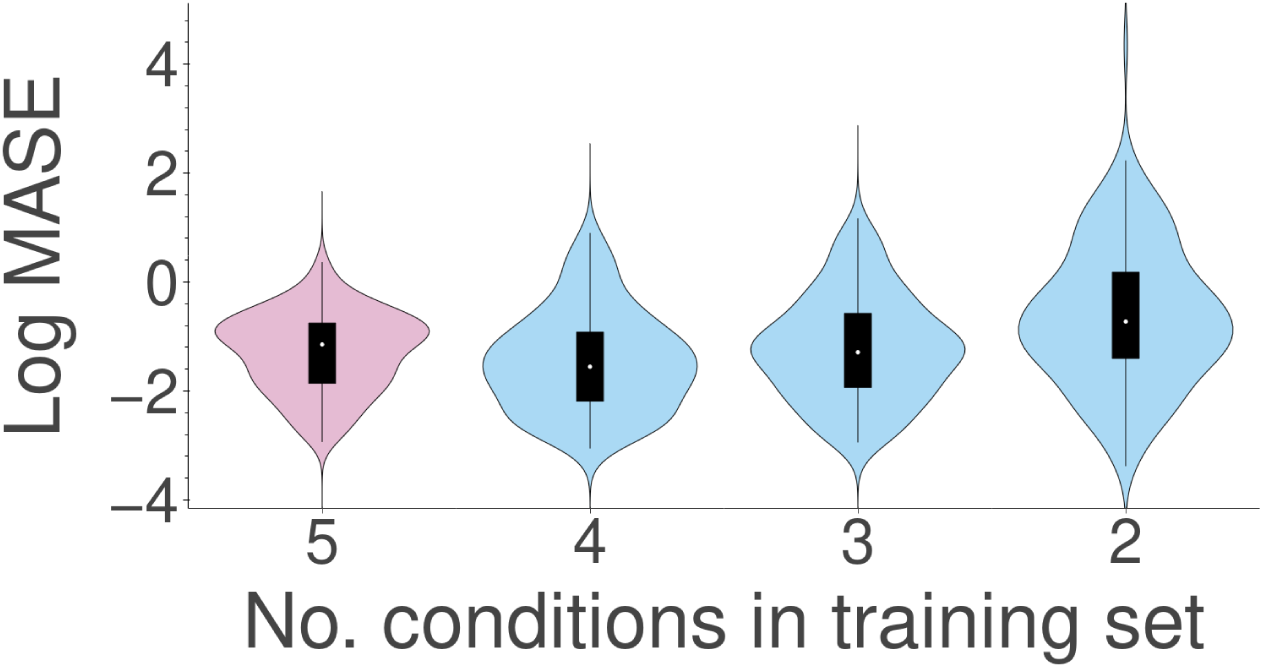
Violin plots of log MASE values corresponding to fine-tuning model shown in Figure 4a on differentially sized training/test sets. The pink (left) violin corresponds to MASE values of all 180 experimental conditions (5 DNA concentrations x 6 3PGA concentrations x 6 Mg^2+^ concentrations); for each of 36 fine-tuning cases, all 5 time trajectories’ data (corresponding to DNA concentrations of 1 nM, 2.5 nM, 5 nM, 7.5 nM, and 10 nM for a particular set of 3PGA and Mg^2+^ concentrations) were provided as training data and as test data. The pink violin is the same as shown in Figure 4b. The blue violins correspond to MASE values where the training and test data were split. In the second violin, for each of the 36 fine-tunings, data corresponding to DNA concentrations of 1 nM, 2.5 nM, 7.5 nM, and 10 nM were provided as training data, and the data corresponding to 5 nM DNA for the same set of 3PGA and Mg^2+^ concentrations were provided as test data. Likewise, in the third violin, for each of the 36 fine-tunings, data corresponding to DNA concentrations of 1 nM, 5 nM, and 10 nM were provided as training data, and the data corresponding to 2.5 nM DNA and 7.5 nM DNA for the same set of 3PGA and Mg^2+^ concentrations were provided as test data. Finally, in the last violin, for each of the 36 fine-tunings, data corresponding to DNA concentrations of 1 nM and 10 nM were provided as training data, and the data corresponding to 2.5 nM DNA, 5 nM DNA, and 7.5 nM DNA for the same set of 3PGA and Mg^2+^ concentrations were provided as test data.

**Figure S4:**
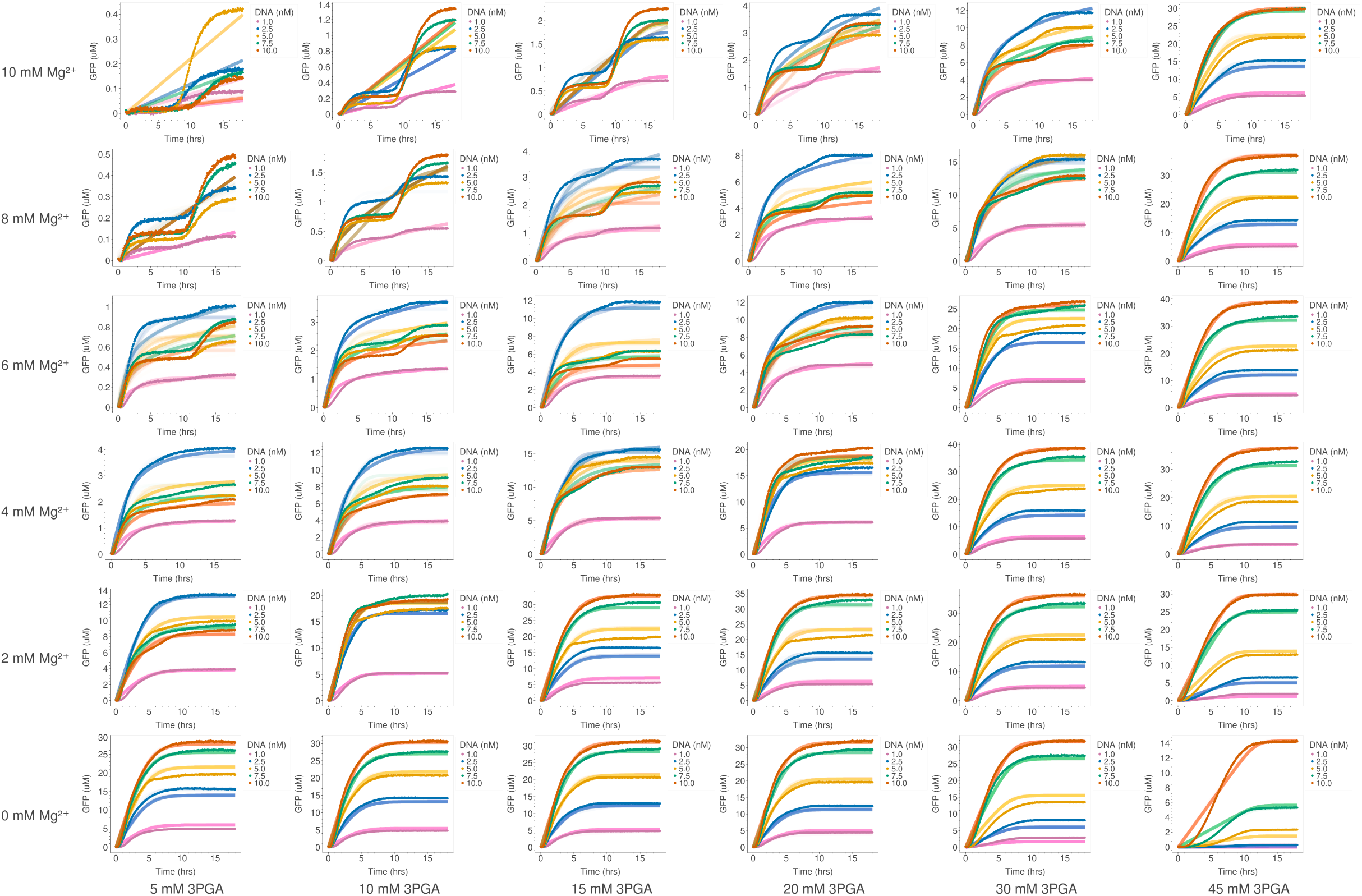
Comparison of model prediction and experimental data after fine-tuning as shown in Figure 4 and using training data as test data. For each of 36 fine-tuning cases, all 5 time trajectories’ data (corresponding to DNA concentrations of 1 nM, 2.5 nM, 5 nM, 7.5 nM, and 10 nM for a particular set of 3PGA and Mg^2+^ concentrations) were provided as training data and as test data. Darker points are experimental data; lighter lines of the same color are model predictions.

**Figure S5:**
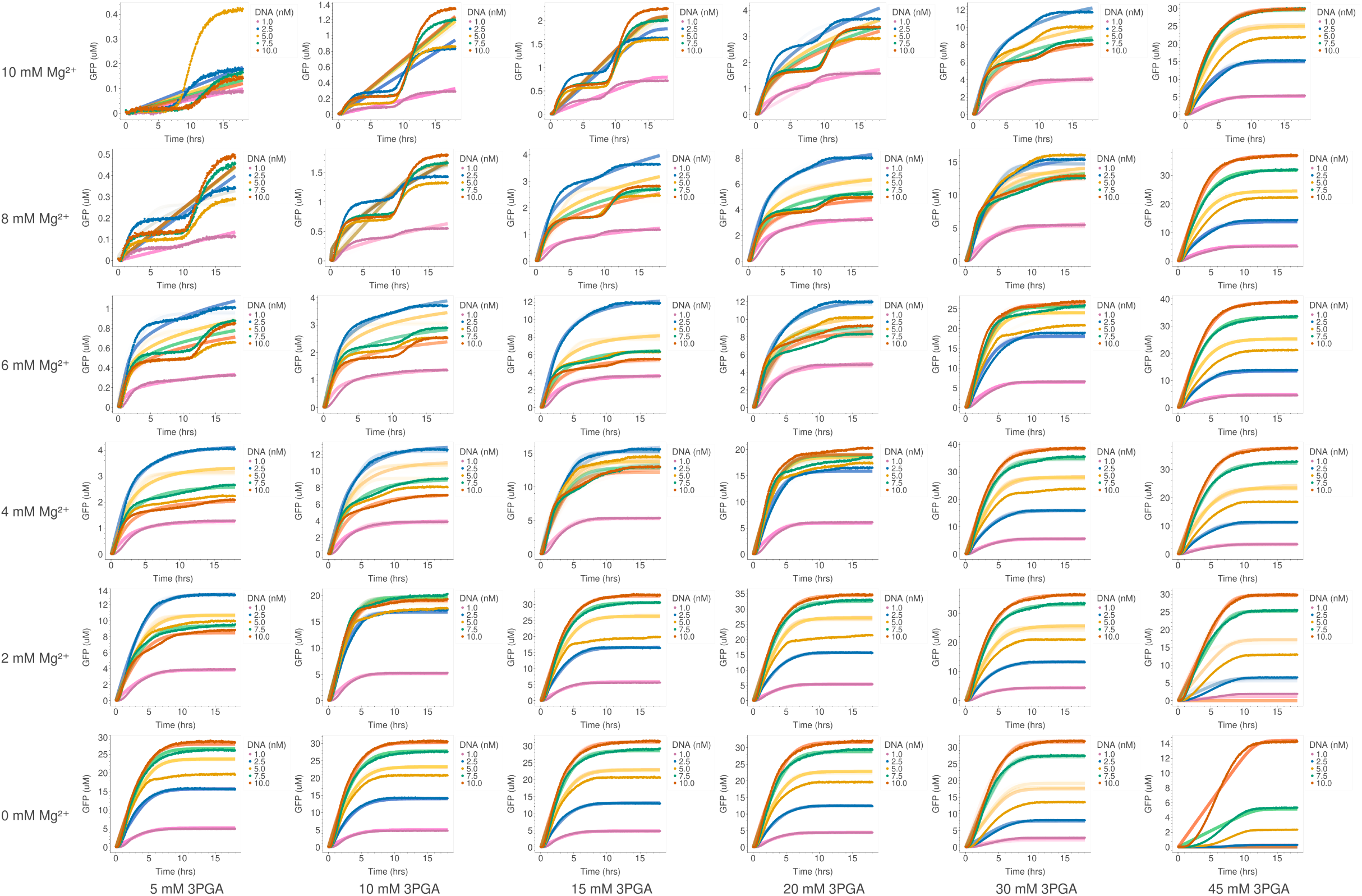
Comparison of model prediction and experimental data after fine-tuning as shown in Figure 4 and using 4 time trajectories as training data. For each of the 36 fine-tunings, data corresponding to DNA concentrations of 1 nM, 2.5 nM, 7.5 nM, and 10 nM were provided as training data, and the data corresponding to 5 nM DNA for the same set of 3PGA and Mg^2+^ concentrations were provided as test data (yellow lines). Darker points are experimental data; lighter lines of the same color are model predictions.

**Figure S6:**
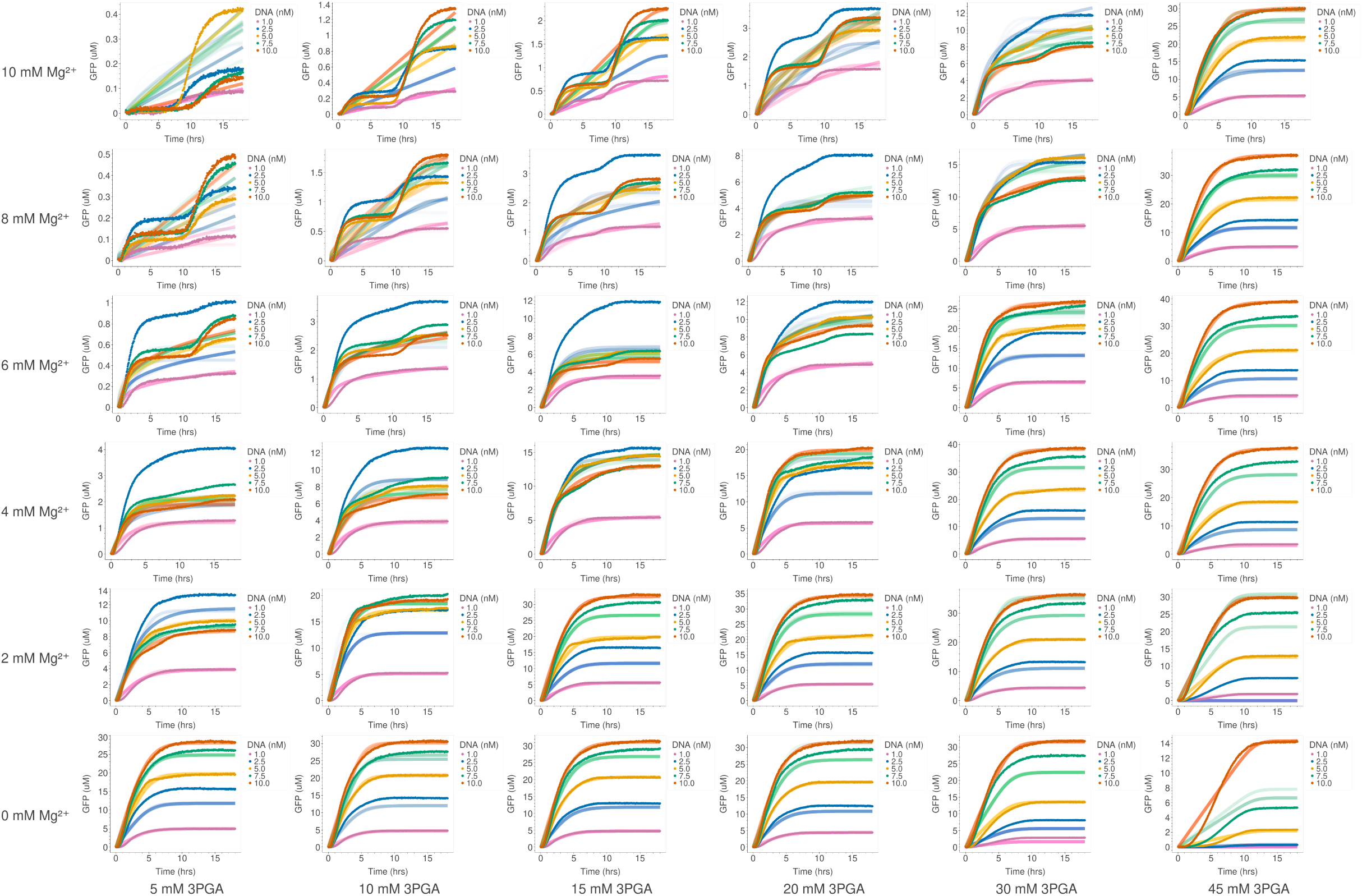
Comparison of model prediction and experimental data after fine-tuning as shown in Figure 4 and using 3 time trajectories as training data. For each of the 36 fine-tunings, data corresponding to DNA concentrations of 1 nM, 5 nM, and 10 nM were provided as training data, and the data corresponding to 2.5 nM DNA and 7.5 nM DNA for the same set of 3PGA and Mg^2+^ concentrations were provided as test data (green and blue lines). Darker points are experimental data; lighter lines of the same color are model predictions.

**Figure S7:**
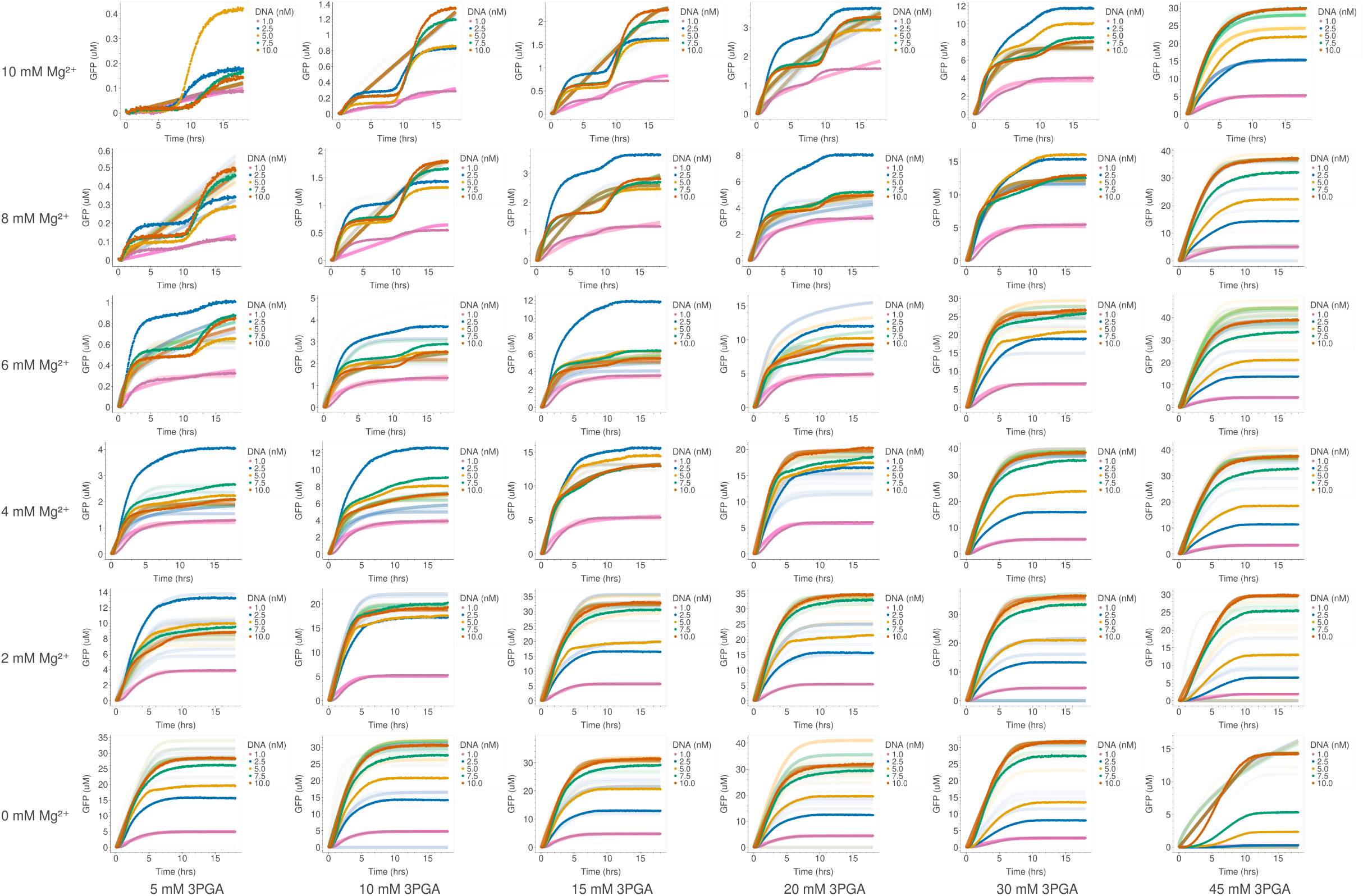
Comparison of model prediction and experimental data after fine-tuning as shown in Figure 4 and using 2 time trajectories as training data. For each of the 36 fine-tunings, data corresponding to DNA concentrations of 1 nM and 10 nM were provided as training data, and the data corresponding to 2.5 nM DNA, 5 nM DNA, and 7.5 nM DNA for the same set of 3PGA and Mg^2+^ concentrations were provided as test data (yellow, green, and blue lines). Darker points are experimental data; lighter lines of the same color are model predictions. Darker points are experimental data; lighter lines of the same color are model predictions.

**Figure S8:**
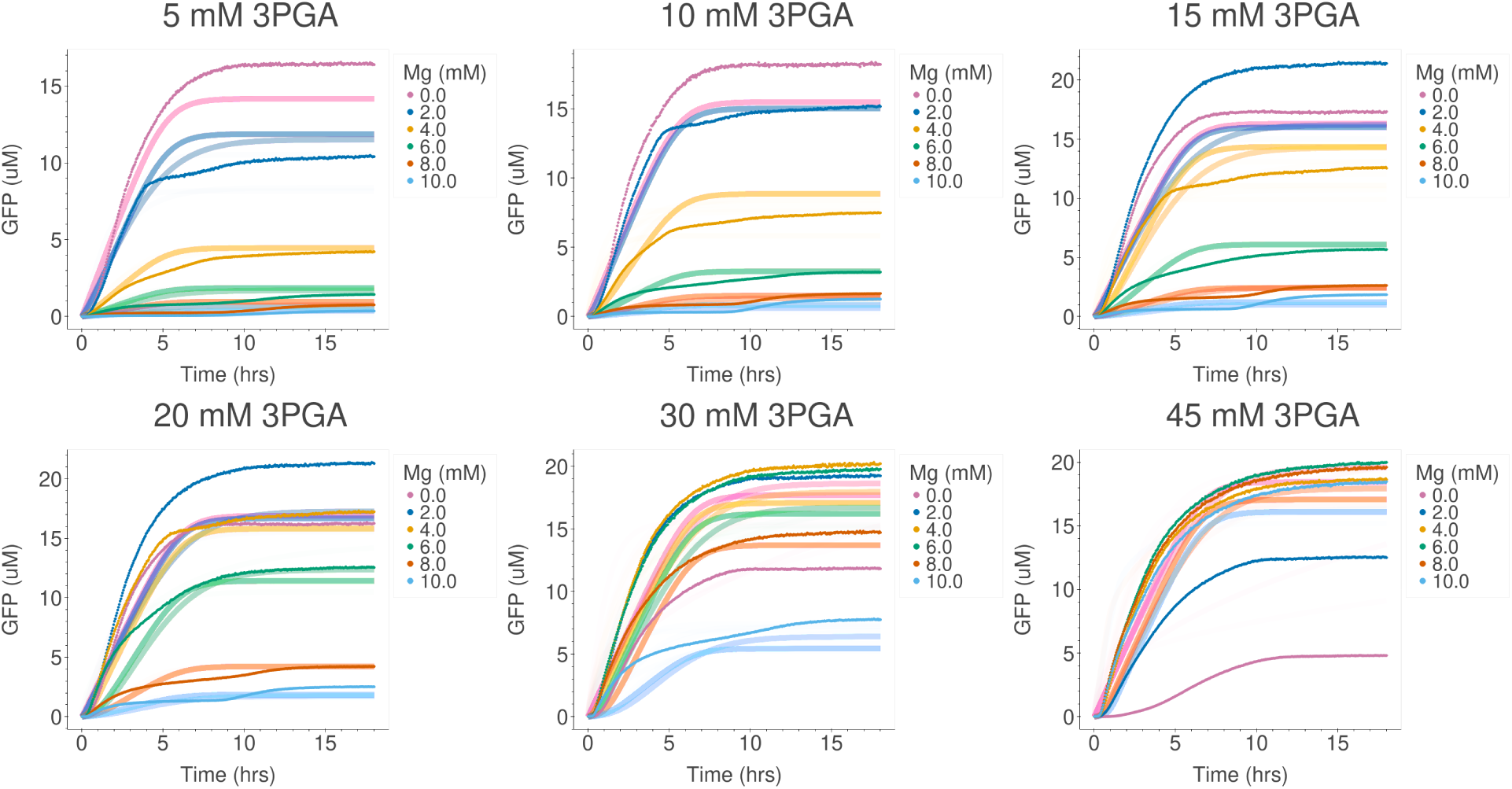
Comparison of model prediction and experimental data after fine-tuning as shown in Figure 5 and using all 36 time trajectories as training data for Lysate Batch 1 from a previously published study [25].

**Figure S9:**
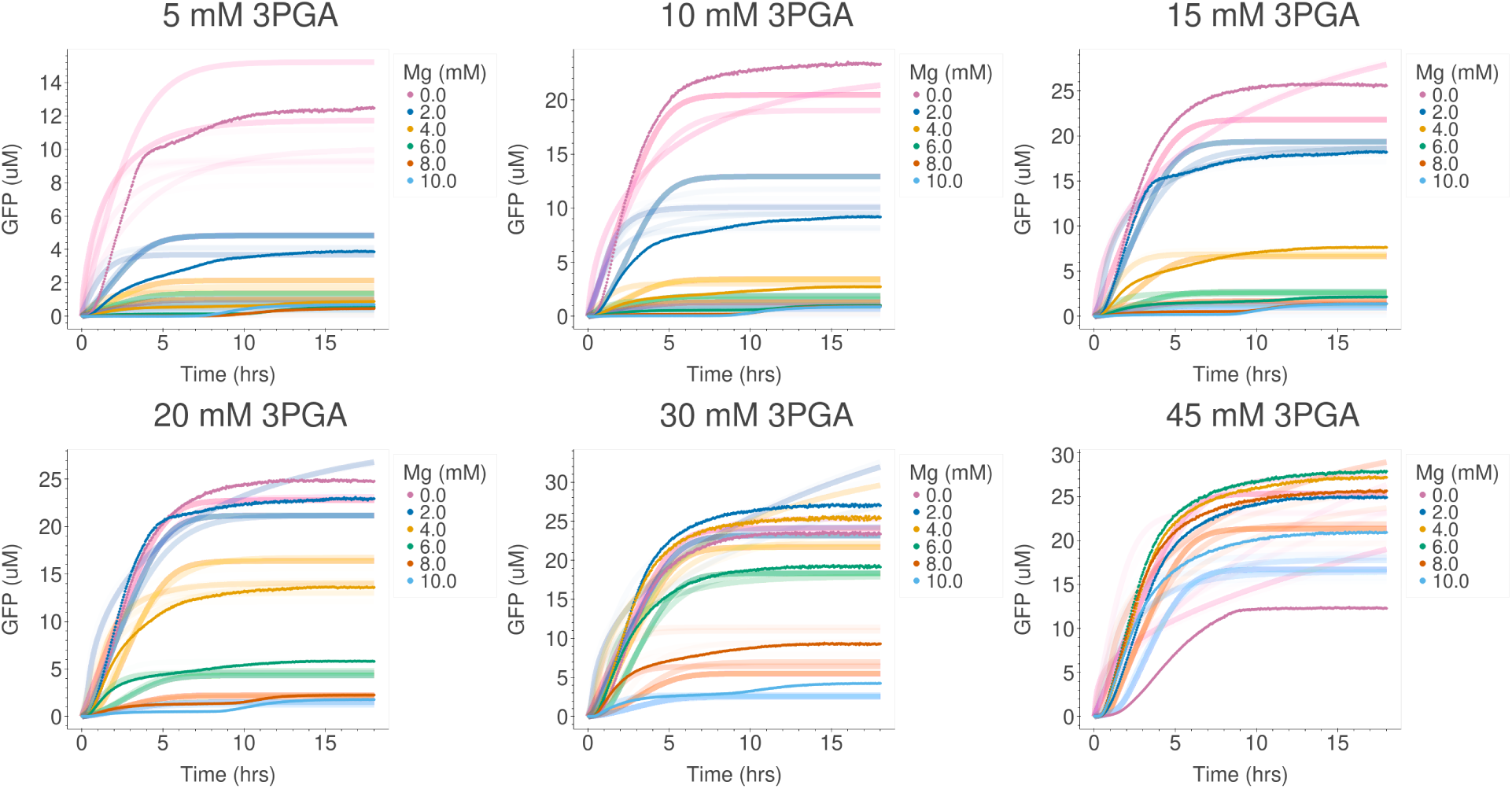
Comparison of model prediction and experimental data after fine-tuning as shown in Figure 5 and using all 36 time trajectories as training data for Lysate Batch 2 from a previously published study [25].

**Figure S10:**
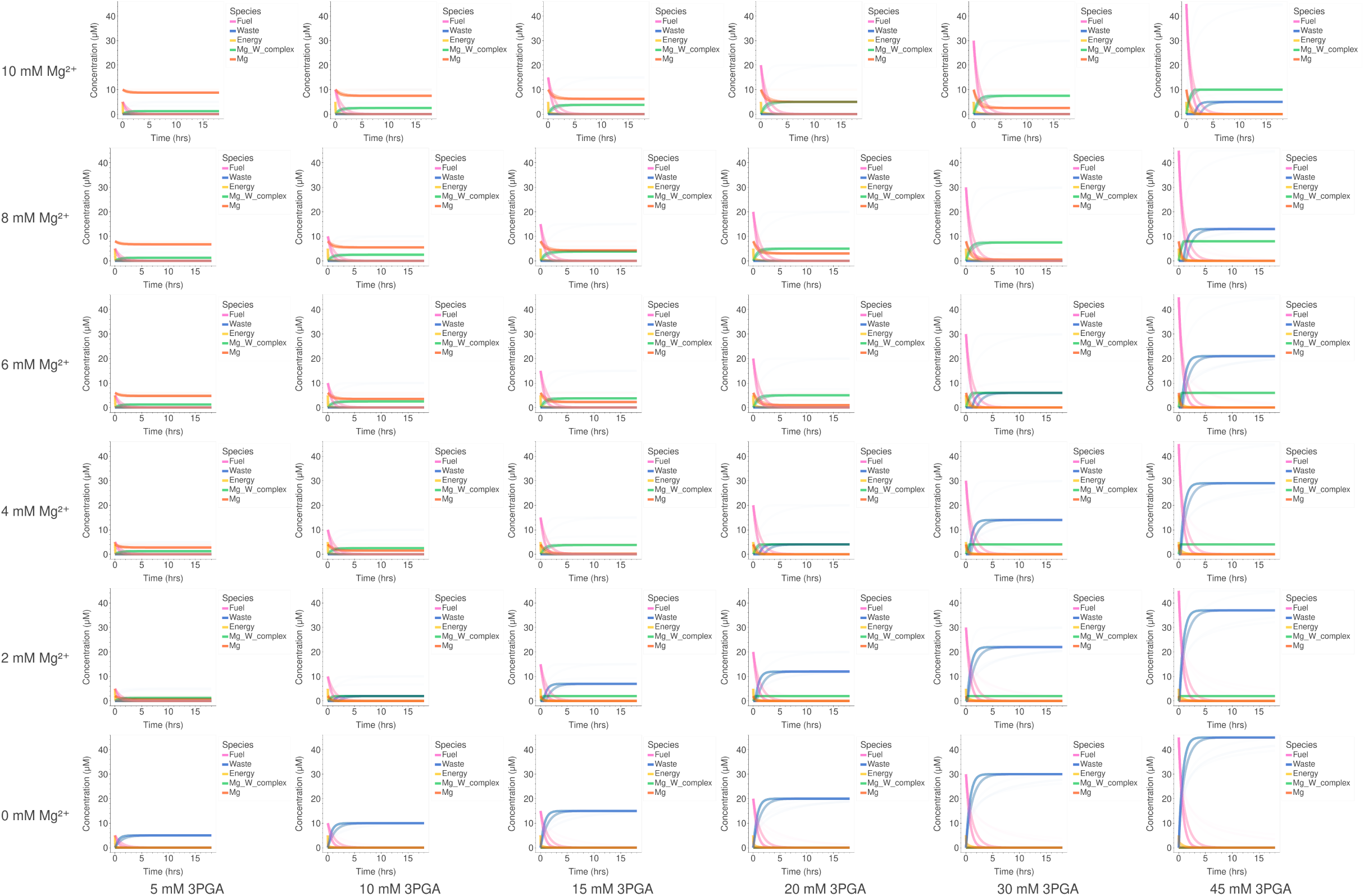
Modeling unmeasured species after fine-tuning as shown in Figure 5 for Lysate Batch 1 from a previously published study [25]. A single model was trained and parameterized for the 36 experimental conditions shown in the plot. The Mg-W complex species is the same as species X in Figure 5A.

**Figure S11:**
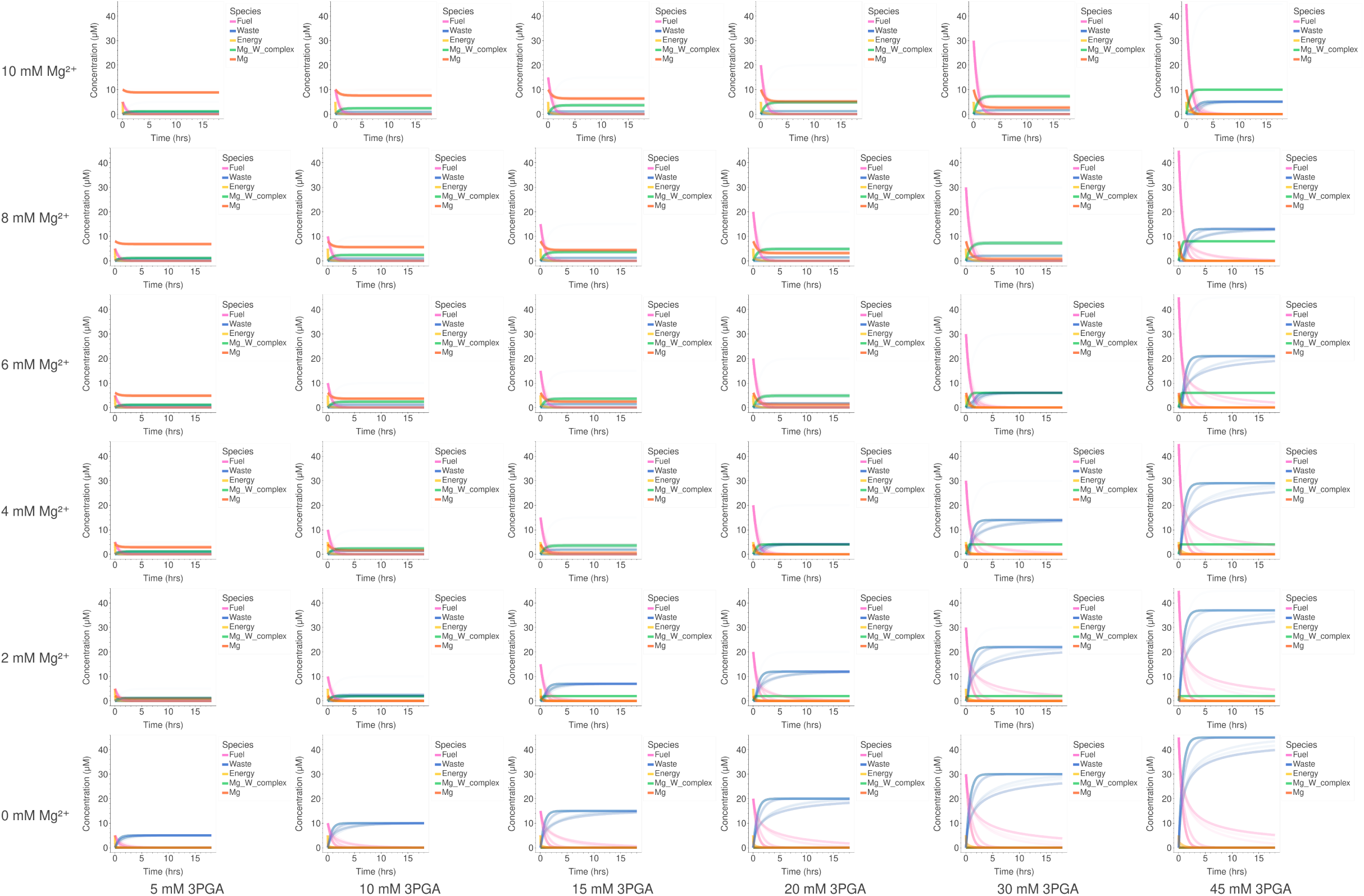
Modeling unmeasured species after fine-tuning as shown in Figure 5 for Lysate Batch 2 from a previously published study [25]. A single model was trained and parameterized for the 36 experimental conditions shown in the plot. The Mg-W complex species is the same as species X in Figure 5A.

